# Murine Abdominal Aortic Aneurysm Intraluminal Thrombus Composition and Structure

**DOI:** 10.1101/2025.10.28.685150

**Authors:** Cortland H. Johns, Luke E. Schepers, Annabelle M. F. Foist, Ethan P. Foster, Anish Bedi, Niharika Narra, Claudia K. Albrecht, Abigail D. Cox, Craig J. Goergen

## Abstract

An abdominal aortic aneurysm (AAA) is a dilation of the aortic wall in the abdomen. Many AAA patients develop intraluminal thrombus (ILT), but the role of ILT in AAA progression and rupture is not well understood. To evaluate ILT in AAAs, we induced AAAs in male C57Bl6/J mice (n=25) via surgical application of topical elastase (5 µL of 5 or 10 mg/mL) to the abdominal aorta below the renal arteries and administration of β-aminopropionitrile (BAPN, 0.2%) drinking water. We collected weekly/biweekly ultrasound images over 56 days. Mice were euthanized and histology images were collected. We semi-quantitatively assessed elastin degradation and inflammation from Movat’s pentachrome and H&E-stained samples, respectively. Mice with ILT had more significant expansion over the length of the study (beginning at day 14, p<0.05). From histology, ILT samples showed more elastin disorganization and greater inflammation. From scanning electron microscopy, we were able to confirm the presence of layered sheets of fibrin and abnormally shaped red blood cells (polyhedrocytes) within the ILT deposits. In this model, elastase causes aortic injury by degrading elastin fibrils in the aortic wall, reducing the ability of the aorta to contract during high-pressure blood flow. Further damage to the extracellular matrix is likely driven by subsequent inflammation. Here we observed tissue samples with greater acute-on-chronic inflammation were correlated with more elastin damage, and therefore greater aortic expansion. Further, larger aortic expansions were correlated with slower blood flow, likely due to increased cross-sectional area. Thus, increased aortic expansion and damage to the aortic wall may be more likely to create hemodynamic conditions that are conducive to the initiation of ILT deposition: endothelial damage and reduced blood flow. Understanding the relationship between ILT formation, aortic wall degradation, and inflammation could help refine therapeutic strategies for treating AAAs.

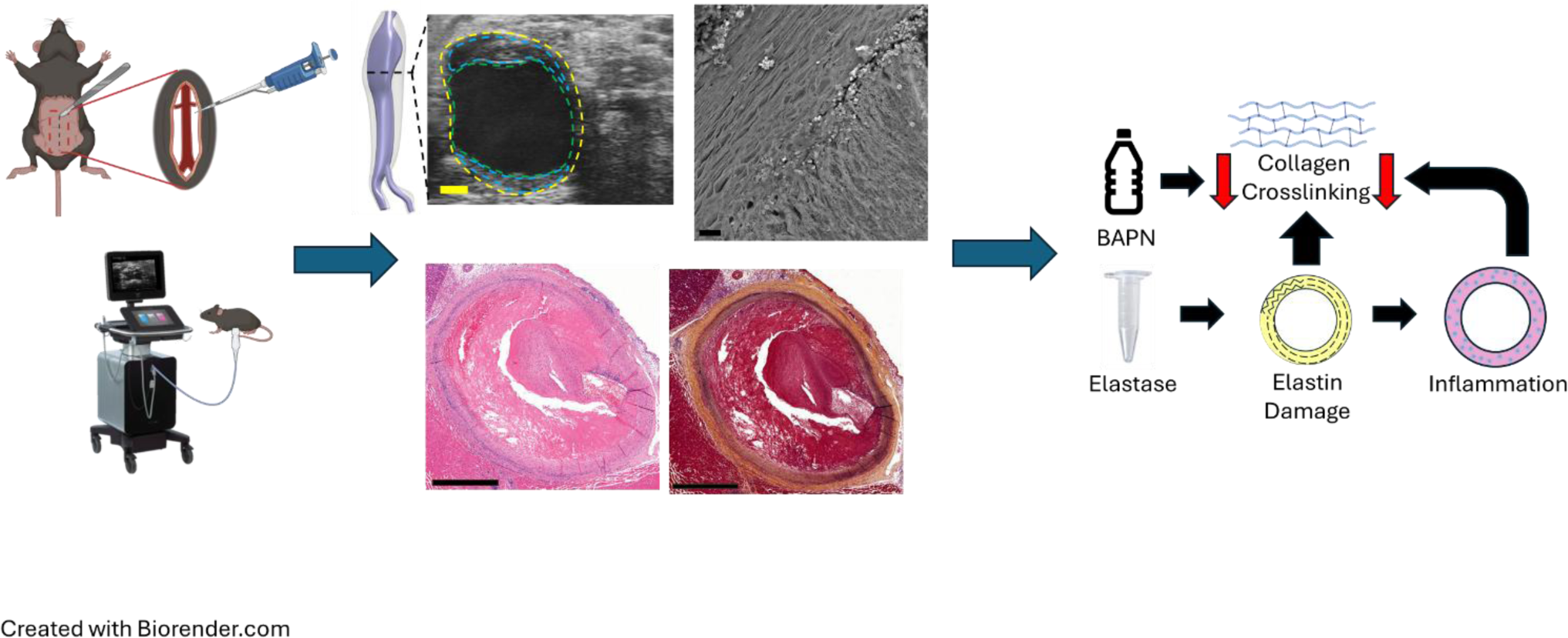

## Introduction

Abdominal aortic aneurysm (AAA), a dilation or ballooning of the aortic wall, frequently occurs near or below the renal arteries (1–4). While some AAAs are asymptomatic and do not increase in size, others continue to enlarge until rupture, frequently causing uncontrolled hemorrhage into the peritoneal space (1, 5, 6). Aneurysm rupture is associated with high mortality rates of 75-90%, resulting in more than 10,000 deaths per year in the United States (1, 7). Even though larger aneurysms are more likely to rupture, there are still instances of smaller aneurysm rupture and larger aneurysm stability (2, 6–8).

Further complicating AAA evaluation in the clinic, 75% of patients with AAAs also develop intraluminal thrombus (ILT) within the expansion (5). ILTs are often heterogeneous and may provide mechanical support to the wall of AAAs, but there are conflicting reports regarding whether ILT increases or decreases stability of expanding AAAs (5, 6, 9–13). In addition to the mechanical role, ILT also serves an active biological function as a locus for the collection of inflammatory cells (8, 10, 13). Thus, a better understanding of how ILT influences aneurysm development and expansion could help improve patient-specific surgical treatment timing and aid in the creation of pharmaceutical treatments of AAA.

Murine disease models provide a repeatable way to evaluate AAA development in a controlled environment on a faster timeline compared to clinical data alone. Several murine models have been established that induce AAAs below the renal arteries using porcine pancreatic elastase (14–16). While one such model induced initial aneurysmal expansion through intraluminal elastase perfusion, these aneurysms did not continue to grow over time, and the occurrence of intraluminal thrombus has not been observed in these models (14, 15). An adaptation of this experimental model includes inducing an aneurysm through topical elastase application to the adventitia, rather than intraluminal application, with the addition of β-aminopropionitrile (BAPN) administration via drinking water (2, 16–19). The administration of BAPN inhibits collagen crosslinking in the adventitial layer during AAA development, allowing for greater expansion than previous models. The topical elastase+BAPN model results in AAAs that can reach 500-600% expansion by 8 weeks post-surgery (18, 19), where higher concentrations of elastase are correlated with increased occurrences of ILT formation (18). Thus, this experimental model may be useful in the study of aneurysmal pathology with a focus on the complex effects of ILT on AAA growth.

Although ILT is present in approximately 75% of human AAA cases, it is unclear whether it serves a protective role by offloading stress from the vessel wall or if it contributes to aneurysm growth and eventual rupture (5, 6, 9–13). In this study, we utilized the elastase+BAPN model described above to evaluate the relationship between aneurysmal expansion, ILT formation, extracellular matrix composition, and inflammation. By combining 4D ultrasound (4DUS), histology, and scanning electron microscopy (SEM), this study provides new insights into how thrombus structure and extracellular matrix remodeling correlate with murine aneurysm expansion. Findings suggest that the presence of ILT is associated with increased AAA severity, suggesting that ILT may be a potential pharmaceutical target to slow the expansion of existing AAAs.

## Results

### All p-values can be found in the supplementary material

#### The topical elastase combined with oral BAPN caused significant and continuous aortic expansion

All of the male mice (n = 23) collectively demonstrated a statistically significant increase in aortic diameter between baseline measurements and each timepoint beginning at Day 14 (*p* ≤ 0.001 for day 14 through day 56 when compared to baseline), as seen in **Figure 2**.

**Figure 1.**
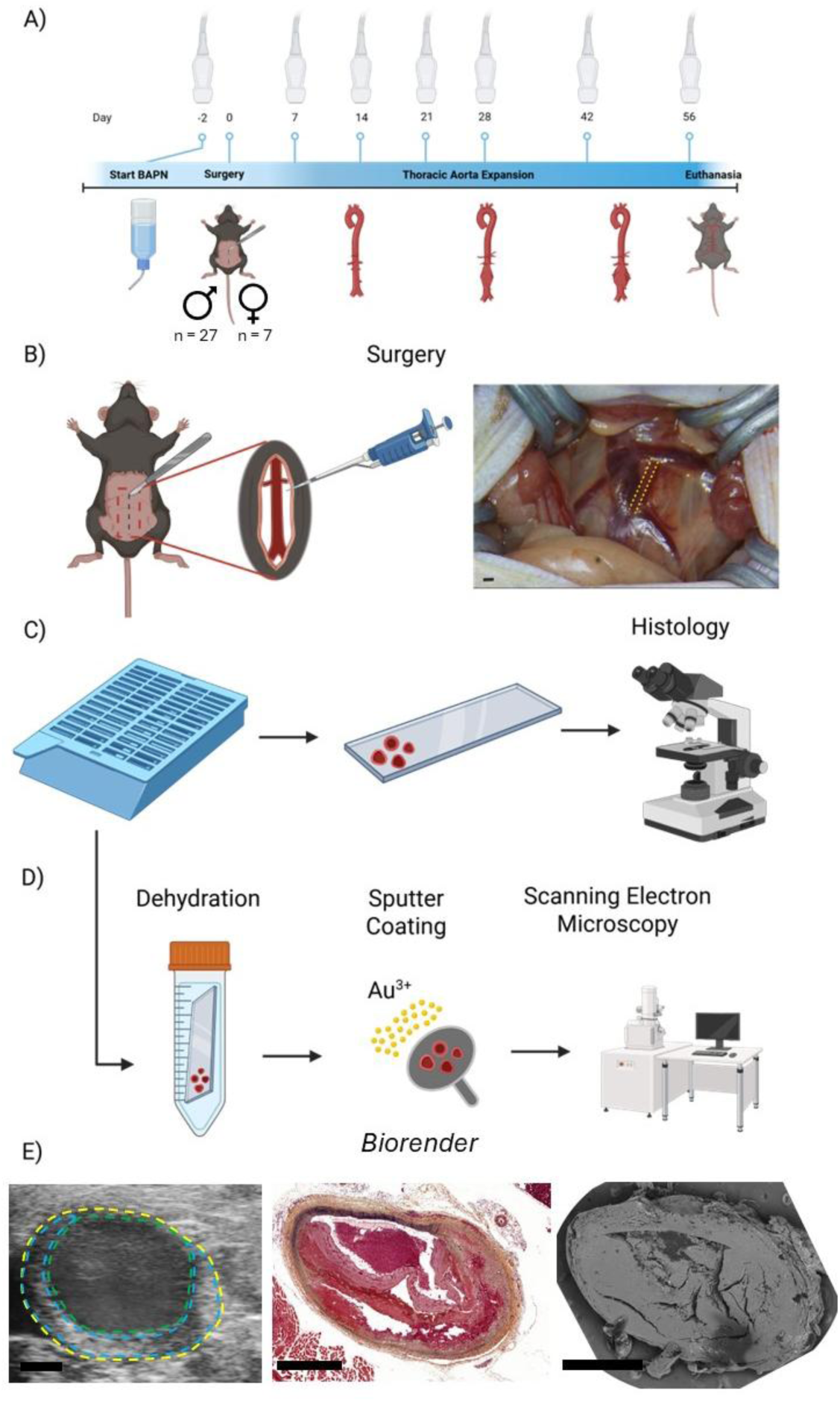
Schematic for the experiments conducted in this study. A) Surgical and imaging timeline for abdominal aortic aneurysm induction in C57Bl/6J male (n = 27) and female (n = 7) mice. B) Application of topical elastase to the aorta (yellow dashed lines) during abdominal aortic aneurysm microsurgeries. Tissue samples from the same paraffin-embedded block can be C) stained for histology or D) dehydrated using increasing concentrations of ethanol and hexamethyldisilazane and sputter coated with a thin layer of gold-palladium to collect scanning electron microscopy. Utilizing the same tissue block leads to decreased dehydration time and better comparisons between scanning electron microscopy and histology. E) We visualized similarity between ultrasound (left), Movat’s pentachrome histology (center), and scanning electron microscopy (right) images. Scale bars: 1 mm.

**Figure 2.**
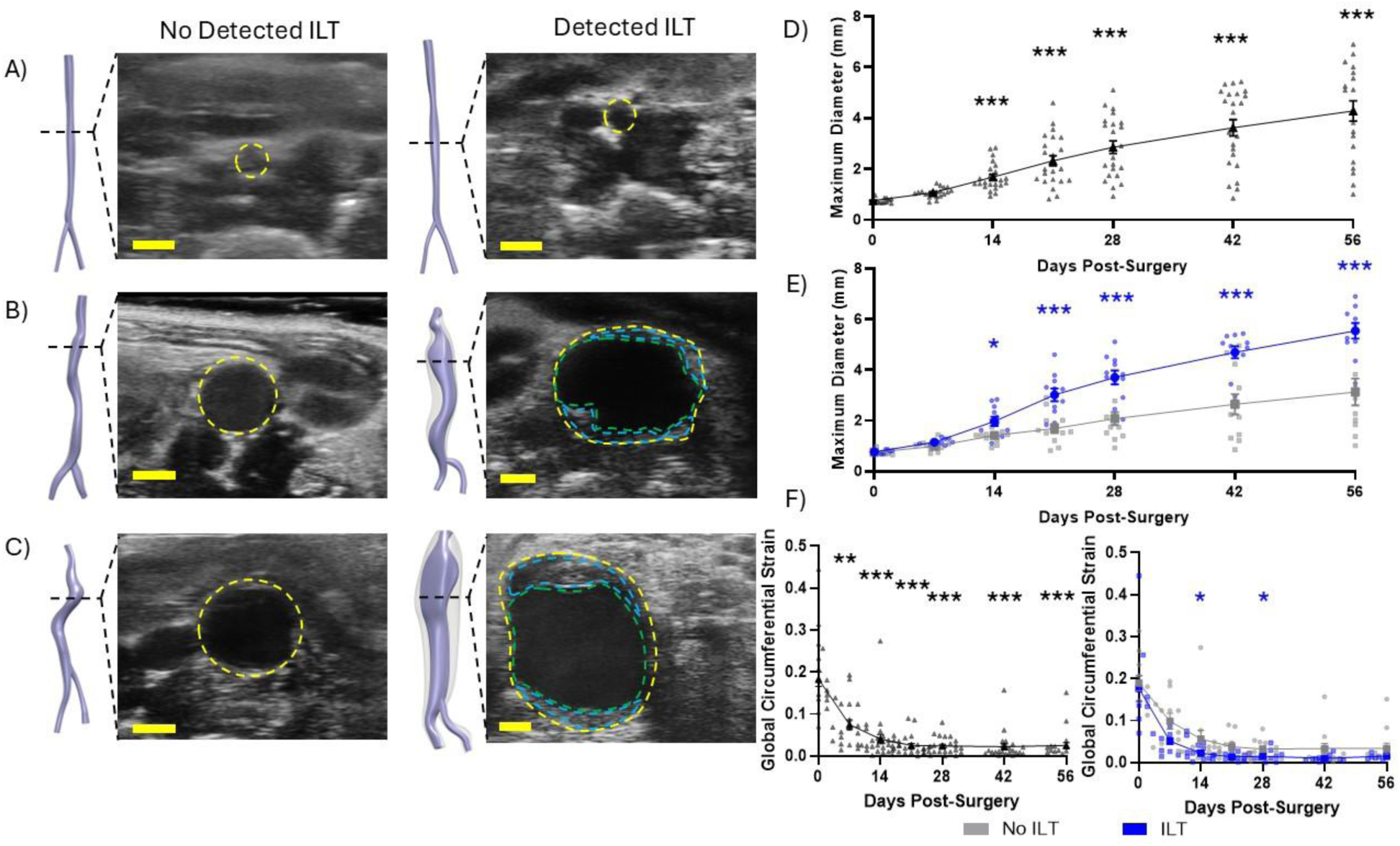
Mice subjected to the topical elastase surgery + oral beta-aminoproprionitrile additive showed continuous growth over time. Short-axis ultrasound images collected from days A) 0, B) 28, and C) 56 from C57Bl/6J male mice are shown with and without intraluminal thrombus (ILT). Vessel wall (yellow dashed), lumen (green dashed) and ILT deposition (blue dashed) highlight this complex pathology. D) Maximum diameter of all male animals over 8 weeks. Statistical significance calculated via a Kruskal-Wallis test and Dunn’s multiple comparison test. ***p≤0.001 - statistically significant difference between baseline and timepoint diameter. E) Diameter of all samples split into detected ILT (blue) and no detected ILT (grey) groups. Statistical significance calculated via a mixed-effects model with the Geisser-Greenhouse correction and Tukey’s multiple comparison test. *p<0.05;***p≤0.001 - statistically significant difference between ILT and no-ILT groups. F) Circumferential strain of all male animals over 8 weeks (left) split into detected ILT (blue) and no detected ILT (grey) groups (right). left: Statistical significance calculated via a lognormal ordinary one-way ANOVA with Tukey’s multiple comparison test. ***p<0.01;***p≤0.001 - statistically significant difference between baseline and timepoint diameter. right: Statistical significance calculated via a lognormal mixed-effects model with the Geisser-Greenhouse correction and Tukey’s multiple comparison test *p<0.05 - statistically significant difference between ILT and no-ILT groups. Bolded data points presented as averages±SE each timepoint. n=12 no ILT; n=11 ILT; scale bars: 1 mm.

#### Samples with detected ILT had greater average expansion and elastin damage than samples without ILT

Of the 23 male mice that survived to the end of the 56-day study, 11 mice developed ILT that was confirmed through histology. There was a significant increase in maximum aortic diameter in samples with ILT compared to those without starting at day 14 and continuing throughout the study as seen in **Figure 2e** (day 14: *p* < 0.05; days 21 through 56: *p* ≤ 0.001). Both groups showed statistically significant differences in aortic diameter compared to baseline beginning at day 7 and remained statistically significant at all timepoints (*p* < 0.05 no ILT; *p* < 0.001 ILT for days 7 through 56 compared to baseline). The group with detected ILT showed significant continuous growth in between each time point (*p* < 0.01 between days 7 and 14; 14 and 21; 21 and 28; and 28 and 42; *p* < 0.05 between days 42 and 56). Between sequential timepoints, the group with no detected ILT only showed significant differences between day 7 and 14 and day 21 and 28 (*p* < 0.05). Further, using the semiquantitative approach to evaluate elastin disorganization (**Figure 3**), all tissue samples with detected ILT had severe elastin damage (n = 11; **Figure 3c-d**). There was greater variability in the elastin damage of tissue samples without ILT (n = 11), with one mouse having no elastin damage, four mice having mild to moderate elastin damage, and six mice having severe elastin damage.

**Figure 3.**
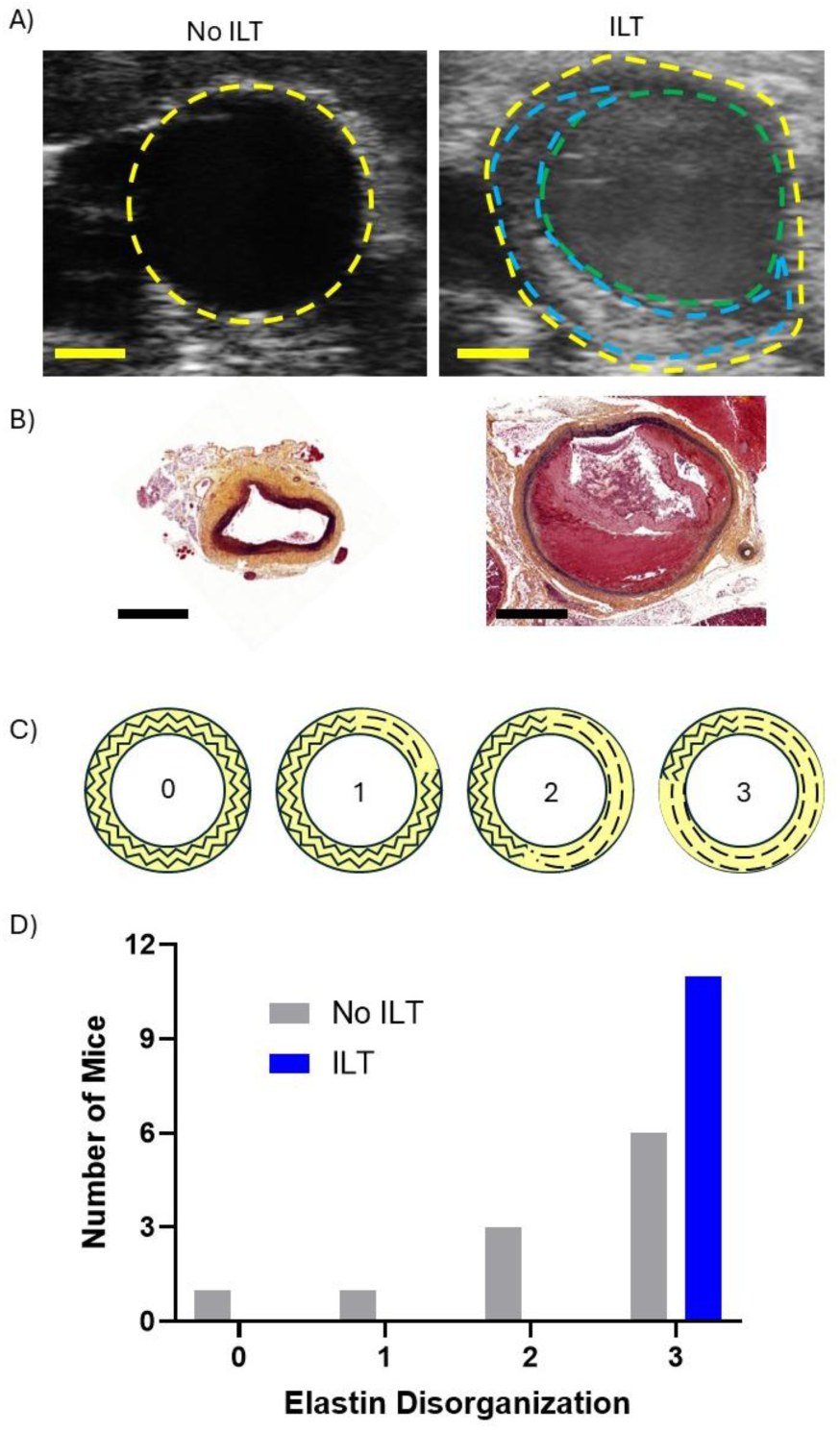
Aneurysms with intraluminal thrombus show more elastin disorganization than samples without observable intraluminal thrombus. A) Short-axis ultrasound images collected from day 56 from samples with and without confirmed intraluminal thrombus (ILT). Vessel wall (yellow dashed), lumen (green dashed) and thrombus deposition (blue dashed) reveal a layered structure. B) Corresponding histology samples stained with Movat’s pentachrome highlight the ILT within the lumen (right). C) Schematic showing degrees of elastin disorder where 0 is normal/no distortion; 1 is less than 25%; 2 is between 25% and 75%; and 3 is greater than 75% of the aortic circumference is affected. D) Semi-quantitative assessment of elastin disorganization. Semi-quantitative scoring determined by a board-certified veterinary histopathologist. n=11 no ILT; n=11 ILT; scale bars: 1 mm.

#### Samples with detected ILT had decreased circumferential strain earlier in the study compared to samples without ILT

All of the male mice collectively demonstrated a statistically significant decrease in circumferential strain between baseline measurements and each timepoint beginning at day 7 (*p* < 0.01 for day 7; *p* ≤ 0.001 for days 14 through 56), as seen in **Figure 2F**. Additionally, there is a significant decrease in strain between day 7 and each remaining timepoint (*p* < 0.05 for day 14; *p* < 0.001 for days 21 through 56). When split into detected ILT and no ILT groups, there was a significant decrease in strain in the ILT group compared to the no ILT group at days 14 and 28 (*p* < 0.05 for all).

#### Extracellular matrix composition differed in the aorta from mice with detected ILT compared to samples without ILT

In samples with detected ILT, the intima showed a substantial increase in the percent composition of collagen compared to the tunica intima of samples without ILT (ILT: 33.9±3.9% collagen; no ILT: 25.9±4.0% collagen; *p* = 0.074; **Figure 4a**). Additionally, there was an increase in the percent composition of proteoglycans in the tunica media in both the ILT and no ILT groups compared to control tissue samples (ILT: *p* < 0.001; no ILT: *p* < 0.01; **Figure 4b**). There was also a considerable increase in proteoglycan content in the ILT group compared to the no ILT group (ILT: 22.8±1.8% proteoglycans; no ILT: 16.9±3.2% proteoglycans; *p* = 0.10). Compared to control aortas, there was a substantial decrease between control and no ILT samples (control: 17.0±2.4% elastin; no ILT: 6.7±1.1% elastin; *p* = 0.052) and a significant decrease between control and ILT samples (ILT: 5.1±0.8% elastin; *p* < 0.05).

**Figure 4.**
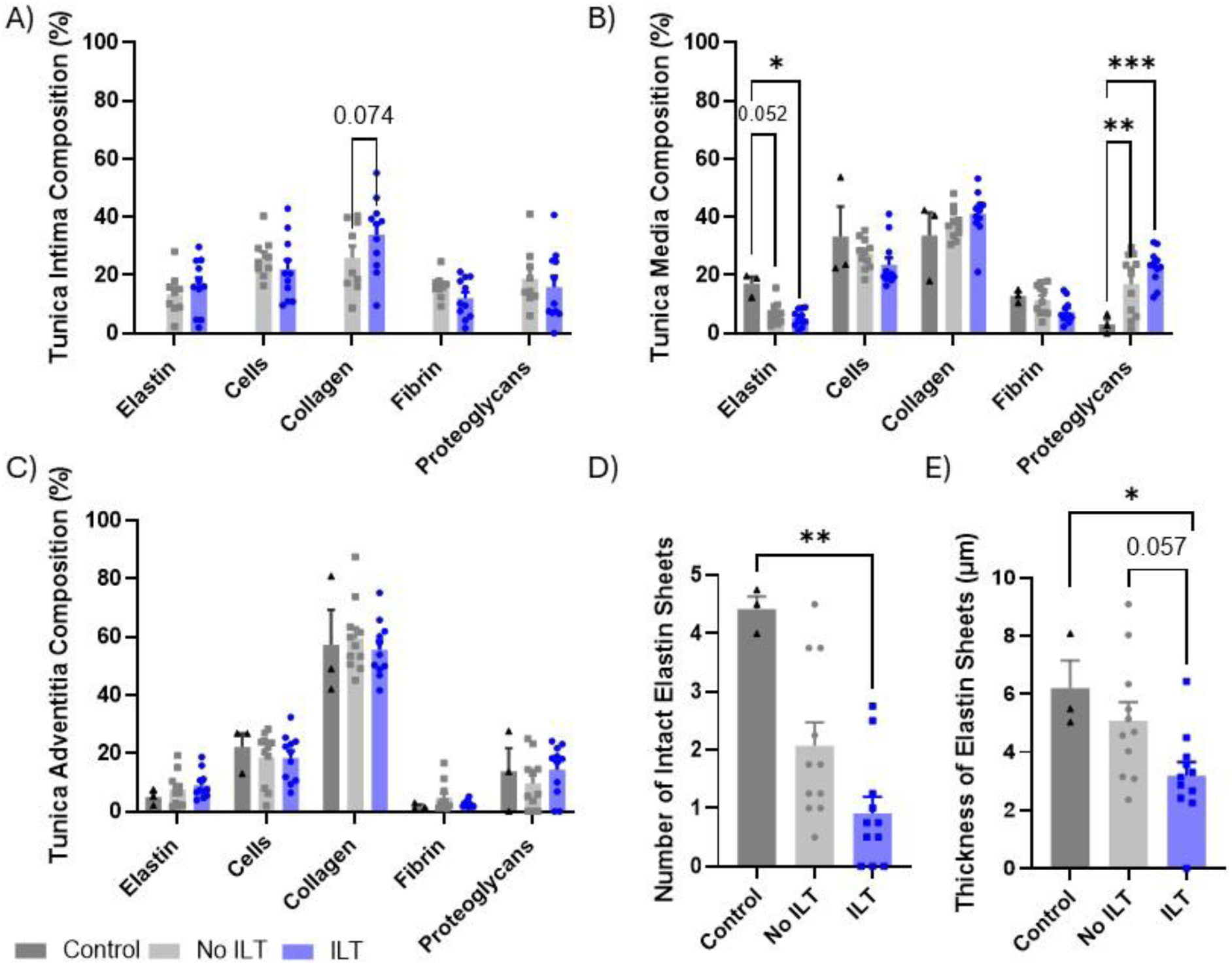
There were distinct changes in extracellular matrix composition at each layer in aortas forming ILT compared to those without confirmed ILT. Extracellular matrix components of the A) tunica adventitia, B) tunica media, and C) tunica intima for each Movat’s pentachrome stained tissue sample were quantified using ImageJ color segmentation. Statistical significance calculated via an ordinary two-way ANOVA with Tukey’s multiple comparison test. Data from tunica adventitia composition underwent a square root transformation to address normality. D) Comparison of the number of intact elastin sheets between samples with and without confirmed ILT, achieved via manual counting of the intact elastin sheets in the wall of the aorta. Statistical significance calculated via a Kruskal-Wallis test with Dunn’s multiple comparison test. E) Comparison of the thickness of the elastin lamellar units between samples with and without confirmed ILT. Statistical significance calculated via an ordinary one-way ANOVA and Tukey’s multiple comparison test. *p<0.05; **p<0.01; ***p<0.001. Bars presented as averages±SE each timepoint. Intima: n=11 no ILT; n=9 ILT; fewer samples were evaluated of the tunica intima due to damage and obscuring of the intima seen in some of the non-thrombus forming samples; control samples were not evaluated for the tunica intima because the healthy intima was too thin to quantify. All other groups n=3 control; n=11 no ILT; n=11 ILT.

There were no significant differences in the extracellular matrix composition of the tunica adventitia between control samples, samples with detected ILT, and samples without (**Figure 4c**). When looking at the number of intact elastin sheets, there was a significant decrease in elastin sheets in samples with ILT compared to control samples (*p* < 0.01), as well as a considerable decrease in the samples with ILT compared to samples without ILT (ILT: 0.9±0.3 sheets; no ILT: 2.1±0.4 sheets; *p* = 0.11; **Figure 4d**). Further, there was a significant decrease in the average thickness of elastin sheets in samples with ILT compared to control samples (*p* < 0.05) and a substantial decrease in the ILT group compared to the no ILT group (ILT: 3.2±0.5 µm; no ILT: 5.1±0.6 µm; *p* = 0.057; **Figure 4e**).

#### Aortas with detected ILT showed greater inflammation than samples without ILT

Tissue samples with detected ILT showed greater inflammatory circumferential involvement, with all ILT samples scoring a 2 or 3 in severity (**Figure 5**). Tissue samples without detected ILT showed more mixed results, with all four severity levels observed across the different tissue samples. (**Figure 5c**). Aortas with ILT trended toward increased polymorphonuclear infiltrates, with all tissue samples but one indicating some level of infiltrate (1 = five mice, 2 = three mice, 3 = two mice), as seen in **Figure 5d**. Samples without ILT, however, had much fewer polymorphonuclear infiltrates, with eight mice having no indication of polymorphonuclear infiltrate. The presence of mononuclear infiltrates was mixed from aortas with and without detected ILT (**Figure 5e**). All samples with ILT had some mononuclear cells present. Over half of the tissue samples with no ILT had no or little mononuclear infiltrate.

**Figure 5.**
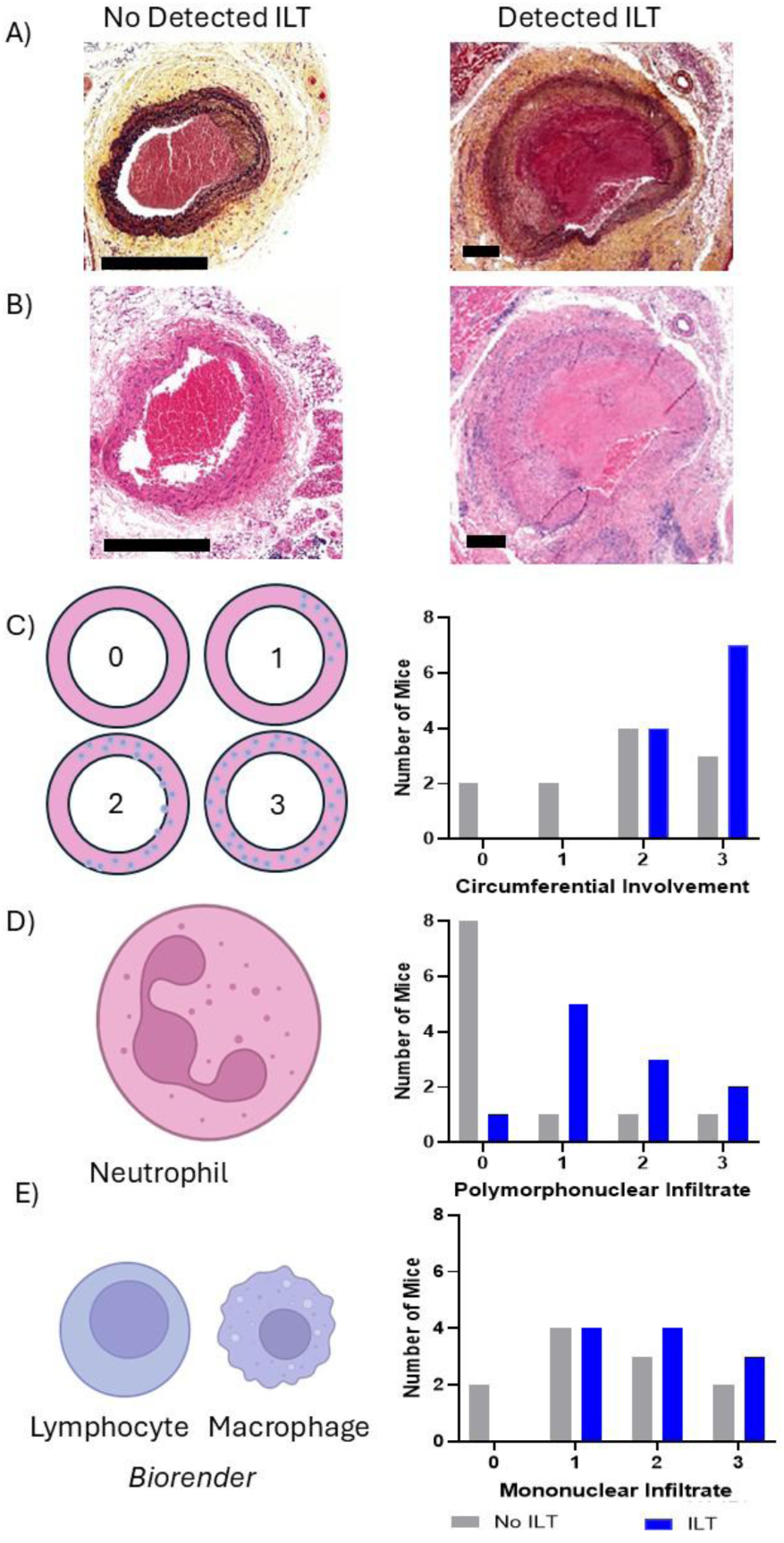
The presence of intraluminal thrombus lead to more acute-on-chronic inflammatory processes. A) Movat’s pentachrome and B) H&E-stained samples of aortic tissue showing minor inflammation (no ILT) and severe inflammation (ILT). Semi-quantitative assessments of C) circumferential involvement of inflammatory cells (% area affected inflammation) where 0 is normal/no inflammation; 1 is less than 25%; 2 is between 25% and 75%; and 3 is greater than 75% of the aortic circumference affected; D) polymorphonuclear infiltrate (acute inflammation) where 0 is normal/no polymorphonuclear infiltrate and 3 is severe polymorphonuclear infiltrate; and E) mononuclear infiltrate (chronic inflammation) where 0 represents normal/no mononuclear infiltrate and 3 is severe mononuclear infiltrate. Semi-quantitative scoring determined by a board-certified veterinary histopathologist. n=11 no ILT; n=11 ILT; scale bars: 300 µm.

#### Scanning electron microscopy revealed RBC transition into polyhedrocytes and disorganized fibrin sheets

Using high-magnification SEM, we acquired additional ILT structure details (**Figure 6**). While we were able to identify the presence of polyhedrocytes within some ILT deposits via scanning electron microscopy (**Figure 6e**), only select ILT had visible red blood cells (RBCs). The majority of thrombus evaluated consisted of fibrin sheets. In some tissue samples, polyhedrocytes were found trapped in these fibrin sheets (**Figure 6g**). Other locations revealed intersecting patterns of fibrin sheets (**Figure 6i**).

**Figure 6.**
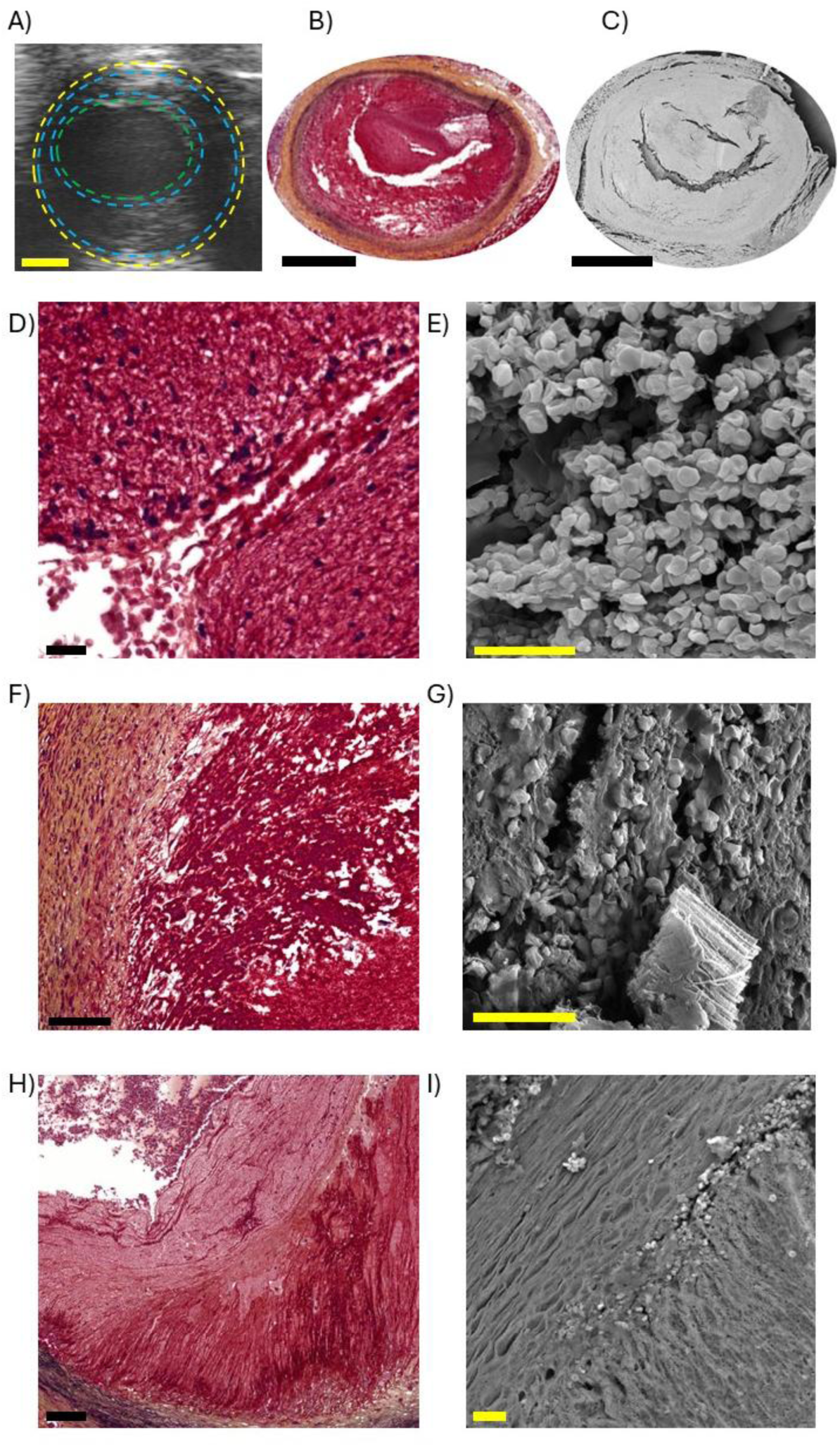
Scanning electron microscopy provided more detail into identifying small structures. Comparison of A) ultrasound, B) Movat’s pentachrome histology, and C) low magnification scanning electron microscopy; scale bars: 1 mm. High magnification images of Movat’s pentchrome histology (D, F, H) and scanning electron microscopy (E, G, I). High magnification SEM examples range from E) polyhedrocyte dense, G) polyhedrocytes trapped in a fibrin mesh (with excess tissue in the bottom left), and I) layered fibrins sheets with different patterns and sparse red blood cells present. Scale bars: 20 µm.

#### Female mice developed more severe and faster growing aneurysms

A much greater proportion of female mice died prior to the end of the study compared to male mice (43% female mortality, 15% male mortality; **Figure 7a**). In comparison to male mice, female animals had significantly larger aortic expansions beginning at day 21 and continuing through day 42 (*p* < 0.05), as well as a considerable increase in diameter compared to males at day 56 (*p* = 0.051; **Figure 7b**). Female mice showed a significant increase in aortic diameter compared to baseline beginning at day 21 and continuing through day 42 (day 21, 28: *p* < 0.05; day 42: *p* < 0.01), with a substantial increase in aortic diameter compared to baseline at day 56 (*p* = 0.053). Additionally, female mice showed significant aortic growth between day 28 and 42 (*p* < 0.05). In the intima, female mice demonstrated substantially greater proteoglycan concentration (*p* = 0.052) and considerably less elastin concentration (*p* = 0.063) compared to male mice. There was similar collagen content compared to male mice with detected ILT (male ILT: 33.9±3.9%, female ILT: 33.5±4.6%), as seen in **Figure 7e**. In the tunica media, female mice had significantly increased proteoglycans compared to combined males as well as males with ILT (combined males: *p* < 0.01; males with ILT: *p* < 0.05), as seen in **Figure 7f** and **Supplemental Figure 1**. Additionally, female mice had significantly lower medial collagen concentrations compared to males with intraluminal thrombus (*p* < 0.05). In the tunica adventitia, female mice had significantly more proteoglycans compared males (*p* < 0.001), as seen in **Figure 7g**. Female mice also had significantly lower adventitial cell concentrations compared to males (*p* < 0.05). In evaluating intact elastin sheets in the media and elastin sheet thickness, female mice with ILT had comparable average values with male mice with ILT. The average intact elastin sheets were 0.9±0.3 sheets for males with ILT and 0.8±0.3 sheets for females with ILT, compared to 2.1±0.4 sheets for males with no ILT, as seen in **Figure 7c**. The average elastin fiber thickness was 3.2±0.5 µm for males with ILT and 3.4±0.9 µm µm for females with ILT, compared to 5.1±0.6 µm for males with no ILT, as seen in **Figure 7d**. From histology grading, we observed that all female mice with ILT had maximal elastin disorganization, as seen in **Figure 7h**. All female mice with ILT were also shown to have maximal circumferential involvement of inflammatory cells, as seen in **Figure 7i**. Additionally, all female mice showed high (>50%) levels of mononuclear infiltrate (**Figure 7k**), but trended toward lower levels of polymorphonuclear infiltrate (**Figure 7j**).

**Figure 7.**
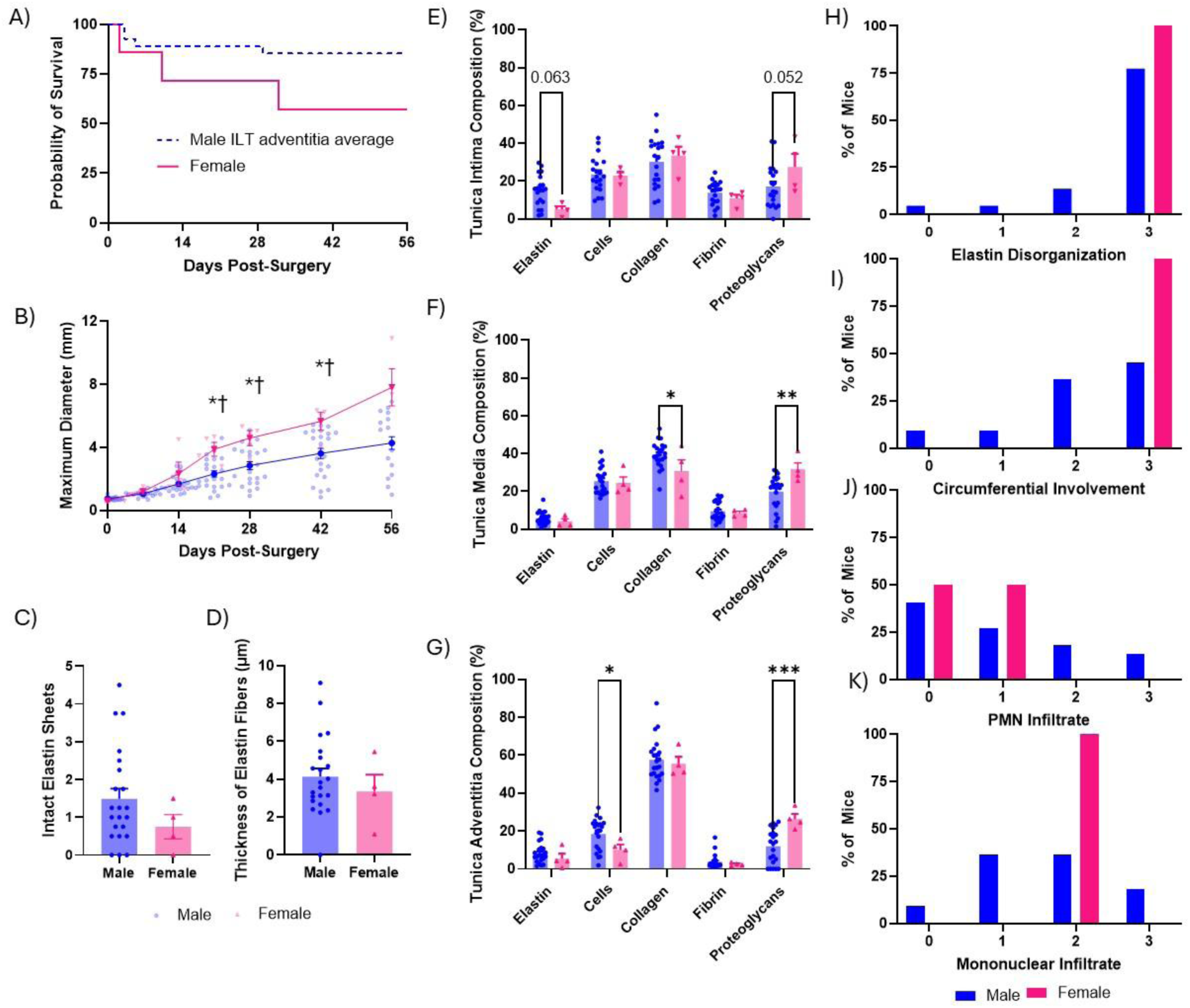
Female mice demonstrated greater response to elastase-BAPN procedure compared to males. A) Number of surviving male and female mice. Final numbers of males included n = 23 with a mortality of n = 4 after surgery. Final numbers of females included n = 4 with a mortality of n = 3 after surgery. B) Diameter of all samples split into female (pink) and male (blue) groups. Bolded data points presented as averages±SE each timepoint. Statistical significance calculated via a mixed-effects model with the Geisser-Greenhouse correction and Tukey’s multiple comparison test. *p<0.05 - statistically significant difference between baseline and timepoint diameter for females; †p<0.05 - statistically significant difference between female and male groups; n = 23 male; n = 4 female. C) Comparison of the number of intact elastin sheets between male and female samples, achieved via manual counting of the intact elastin sheets in the wall of the aorta. Statistical significance calculated via a Mann-Whitney test. D) Comparison of the thickness of the elastin lamellar units between male and female samples. Statistical significance calculated via a Welch’s t-test. Extracellular matrix components of the E) tunica adventitia, F) tunica media, and G) tunica intima for each Movat’s pentachrome stained tissue sample were quantified using ImageJ color segmentation. Statistical significance calculated via an ordinary two-way ANOVA with Tukey’s multiple comparison test. *p<0.05;**p<0.01; ***p<0.001. Bars presented as averages±SE each timepoint. Semi-quantitative assessments of H) elastin disorganization where 0 is normal/no distortion; 1 is less than 25%; 2 is between 25% and 75%; and 3 is greater than 75% of the aortic circumference is affected; I) circumferential involvement of inflammatory cells (% area affected inflammation) where 0 is normal/no inflammation; 1 is less than 25%; 2 is between 25% and 75%; and 3 is greater than 75% of the aortic circumference affected; J) polymorphonuclear infiltrate (acute inflammation) where 0 is normal/no polymorphonuclear infiltrate and 3 is severe polymorphonuclear infiltrate; and K) mononuclear infiltrate (chronic inflammation) where 0 represents normal/no mononuclear infiltrate and 3 is severe mononuclear infiltrate. Semi-quantitative scoring determined by a board-certified veterinary histopathologist. Intima: n=20 male; n=4 female; fewer samples were evaluated of the tunica intima due to damage and obscuring of the intima seen in some of the non-thrombus forming samples. All other groups n=22 male; n=4 female.

## Discussion

In this study, we induced murine AAAs with a combination of topical elastase and BAPN. We observed continuous AAA growth over eight weeks with 48% of males and 100% of females developing ILT. Aneurysms that developed ILT were associated with larger maximum diameters, greater presence of inflammatory cells, and significant differences in extracellular matrix composition. ILT samples consisted primarily of thick sheets of fibrin with some formation of polyhedrocytes.

With an average expansion of greater than five times the initial diameter at day 56 (baseline: 0.76±0.02 mm; day 56: 4.3±0.4 mm; **Figure 2d**), results were comparable to other studies that have utilized this murine model (16–19). In the acute stage during and immediately after surgery, damage to the aorta was initiated by exposing the vessel, removing the surrounding connective tissue, and topically applying elastase to the region ranging from the renal arteries to the iliac bifurcation (17, 18). Elastase caused injury to the aorta by degrading the elastin fibrils present in the wall of the aorta and inducing an inflammatory response, reducing the ability of the aorta to store elastic potential energy (2, 18–21). After this degradation, the body cannot produce more functional elastin, so the healing mechanism was dominated by the deposition and remodeling of collagen in the elastase-deficient area, strengthening the extracellular matrix through fiber crosslinking (3, 16, 18). Our data support the findings that the use of BAPN, a lysyl oxidase inhibitor, inhibits crosslinking, causing a secondary insult that leads to continued expansion (17, 19, 20, 22–24). Thus, the continued AAA growth over time emulated the many aspects of the continued progression of aortic aneurysm growth in humans (18, 19). Both the continuous infrarenal growth and the varied presence of ILT emulated much of the human pathology, thus making this an effective comparative model to investigate the role of ILT in human AAA disease. While previous studies have utilized this model to evaluate AAA growth (16–19), we focused this study on evaluating the relationship between ILT, ECM components, and inflammation.

Since this elastase+BAPN model directly targeted elastin, the decreased elastin in the tunica media in treated mice (both with and without ILT formation) compared to healthy tissue was expected (Figure 4), as previous studies of human infrarenal AAAs also found decreased elastin content (4, 22, 25, 26). This confirms that the applied elastase worked properly to damage elastin present in the aortic wall. This damage is further supported by elastin degradation grading done through histology, where all male mice except one showed at least partial damage to elastin. In addition to decreased elastin, we also found increased proteoglycans of both the ILT and no ILT forming groups compared to control aortic tissue (Figure 4). Increased proteoglycan content is a common indicator of aortic medial damage in aortic aneurysms (27–30). This further supports the idea that this animal model induces aneurysms that mimic key aspects of human pathology. Further, previous studies have established that a healthy mouse aorta typically has 5 elastin lamellar units (31, 32). In control mice, we found an average of 4.4±0.2 elastin sheets, suggesting that our method of counting elastin sheets aligns well with previous findings.

We found that AAAs with detected ILT had larger expansions compared to AAAs without thrombus (**Figure 2**). A similar correlation between AAA diameter and ILT volume has been reported in previous clinical studies (11, 12), where more severe aneurysm expansion was associated with ILT formation. This suggests that intraluminal thrombus may be associated with a worse prognosis for AAA expansion. That said, it is not clear if ILT formed because there was a greater expansion to begin with and provided the hemodynamic conditions for ILT deposition, or if ILT drove the greater expansion in the first place. In addition, a significant decrease in strain after elastase treatment was expected, as damaged elastin causes the arterial wall to stiffen (33–35). However, the ILT group showed greater decreased strain compared to aortas without ILT at earlier timepoints in the study. This suggests that AAAs with ILT are subjected to greater stiffening and less wall motion at an earlier phase than seen in AAAs without ILT, with both groups ultimately approaching nearly no wall motion by the end of the study. Histological analysis also found several interesting distinctions in the intima and media between murine AAAs with ILT and those without. The greater concentration of collagen in the intima from mice that developed ILT is an indicator of intimal hyperplasia, a pathological buildup due to vascular smooth muscle cell migration and inflammatory cell infiltration (36–46). Previous studies have shown similar intimal hyperplasia (47, 48), suggesting that this is not a finding specific to the elastase+BAPN model. Additionally, samples with ILT showed greater proteoglycan concentrations in the tunica media compared to samples without ILT, indicating greater damage to the media in ILT-forming samples. These findings of increased aortic medial and intimal remodeling in AAAs with ILT suggest that AAAs with ILT demonstrate greater damage to the aortic wall, and therefore more severe disease progression.

While the decrease in elastin content was modest between samples with and without ILT, we observed fewer intact elastin sheets, decreased average elastin fiber thickness, and increased elastin deformation in AAAs with ILT. Similar elastic fragmentation has been seen in other animal models utilizing porcine pancreatic elastase as well as human studies of AAAs (24, 26, 47), and elastin fragmentation has been shown to lead to proteoglycan accumulation (27). This indicates that while similar concentrations of elastin may still have been present in the media of AAAs with and without ILT, elastin is less functional in samples with ILT, which is correlated with greater AAA growth in ILT-forming samples. Additional preclinical and clinical AAA studies characterizing elastin function, degradation, and preservation strategies are warranted.

We also found that AAAs with ILT showed greater indicators of inflammation including circumferential involvement, polymorphonuclear infiltrate, and mononuclear infiltrate (**Figure 5**). Notably, very little polymorphonuclear infiltrate was detected in samples without ILT (**Figure 5**), indicating a reduced level of acute inflammation in these AAAs lacking thrombus. This suggests that greater acute-on-chronic inflammation may be present in AAAs with ILT compared to those without, providing evidence for a connection between inflammation and thrombus formation. This finding is supported by a previous study that found greater inflammatory cell content in human AAAs with larger samples of ILT compared to those with little or no ILT (8).

The observation that murine aortas with ILT also had increased average maximum diameter, greater damage to elastin sheets, and more severe inflammatory processes suggests that the presence of ILT may worsen AAA expansion through an increased inflammatory response. While elastase caused initial injury to the aorta by degrading the elastin fibrils and sheets in the aortic wall, further damage to the extracellular matrix within the wall is often driven by subsequent acute and chronic inflammation (49). Here we observed that the tissue samples displaying greater acute-on-chronic inflammation are correlated with more damage to elastin, likely allowing for greater expansion of the aorta. Further, larger aortic expansion was correlated with slower blood flow velocity due to the increased cross-sectional area of the vessel (**Supplement 2**). Increased aortic expansion and damage to the aortic wall may be more likely to allow for the hemodynamic conditions that are conducive to the initiation of ILT deposition: damage to the aortic endothelium, decreased wall shear stress, and increased particle residence time (13). Understanding the relationship between ILT formation, aortic wall degradation, and inflammation could help refine clinical strategies for monitoring and treating AAAs. Based on this understanding, potential treatments to slow the progression of AAA development could include platelet-targeting or anti-inflammatory therapies that localize to this region of the systemic vasculature. This strategy is supported by a previous study evaluating infrarenal AAAs in human patients taking antiplatelet medications prior to AAA repair surgery (50). This clinical study found that patients with AAAs taking aspirin had lower ILT area and higher concentrations of arterial elastin (50). Future work focused on modulating inflammation and ILT formation could offer promising pathways for non-surgical intervention in AAA treatment.

SEM can be used to visualize the structural components of biological tissue with unrivaled detail. In a previous study utilizing a murine aortic dissection model, SEM was used to visualize the structural components of intramural thrombus (IMT) revealing abnormal, polyhedral-shaped RBCs and significant fibrin after IMT maturation and contraction (51). While the structure of RBCs in IMT played an important role (51), we observed in this study that polyhedrocytes were present in only in some samples (8/20 regions of interest (ROIs)). We more commonly found large concentrations of fibrin (19/20 ROIs). This indicates that there was a substantial difference in thrombus structure in AAAs compared to aortic dissections, likely due to hemodynamic differences. While blood flow was slowed in expanded AAAs, blood was still moving in the lumen, compared to relatively stagnant blood pooled in the false lumen of an aortic dissection. This could suggest that RBCs are likely to become trapped at a much higher rate in the false lumen of an aortic dissection compared to the ILT in an AAA. Indeed, fibrin structures may be necessary in regions where active flow is present in order to stabilize the thrombus. This is supported by the previous study of IMT, where fibrin was most prevalent in the aortic region closest to the false lumen opening (51).

Finally, we observed clear sex-specific differences in AAA development. Female mice displayed significantly greater mortality compared to males, indicating a heightened sensitivity to the combined insult of elastase+BAPN. Clinical data has shown that while AAA occurrence is more common in men, women with AAAs have a higher likelihood of rupture and higher mortality rates (52–57). Thus, this model emulates sex differences seen in clinical settings. Among the female surviving mice (n=4), these mice demonstrated larger aneurysmal expansions beginning at Day 21 and continuing throughout the study compared to male mice (**Figure 7**). This indicates a more severe disease progression in female mice compared to males, further aligning with existing clinical data (52–57). Other histological indicators of disease severity, including elevated proteoglycan accumulation across all layers of the aortic wall and decreased elastin concentration in the tunica media compared to males were also different between male and female mice. Interestingly, female animals also had lower concentrations of collagen in the tunica media as well as lower concentrations of cells in the tunica adventitia compared to male mice. While these findings could be due to the low sample sizes of females utilized in this study, they could also be due to decreased ECM components and subsequent decreased mechanical support of the expanded aorta in female animals (Figure 7). These sex-specific differences could explain the greater likelihood of AAA expansion and rupture seen in females compared to males. Histological assessment of inflammation demonstrated more pronounced circumferential involvement and mononuclear infiltrates in female mice compared to males. These findings indicate an even greater sustained chronic inflammatory processes compared to male animals. However, female mice trended towards lower levels of acute inflammation. This could be an indicator that new inflammatory processes are occurring less frequently in female mice, with severe aortic expansion being primarily driven by inflammation earlier in the disease process, rather than the continued, ongoing damage seen in males, but additional studies are needed to fully elucidate inflammation timing. Cumulatively, these results support that females experience more severe AAA progression characterized by higher mortality, greater aneurysmal growth, enhanced proteoglycan deposition, and amplified chronic inflammatory responses compared to males. Since previous studies have found that aneurysms rupture in females at smaller sizes (54, 55, 57), and women are less likely to be recommended for surgery to repair an AAA (52–57), this could be an indicator that women should be considered for AAA repair surgery sooner in the disease process than men (52, 53).

This study has several limitations. First, the number of animals that survived until the end of study was relatively small. However, this limitation is mitigated by the longitudinal design of the study, in which multiple imaging timepoints provided a large dataset for tracking disease progression, and the fact that statistically significant results were observed from both the ultrasound and histological analysis. Second, while three metrics of inflammation were evaluated, additional histological and molecular markers could provide a more comprehensive picture of inflammatory processes in AAA development. Incorporating a broader range of inflammatory assessments in future studies could help identify underlying molecular mechanisms. Finally, although both male and female mice were included in this study, additional work focused on sex-differences specifically powered to address these differences could help confirm the trends we observed to fully characterize sex-specific influences on AAA progression.

In conclusion, elastase+BAPN murine aneurysms with ILT were larger, stiffer, and exhibited greater collagen and proteoglycan deposition and inflammatory infiltration compared to those without ILT. Clinically, these findings suggest ILT formation correlates with more severe disease. The observed link between ILT and inflammation supports anti-inflammatory or platelet-targeted therapies as potential non-surgical interventions, consistent with human data showing smaller ILT burden in aspirin users. Sex differences were pronounced, with females exhibiting higher mortality, faster expansion, and stronger chronic inflammation. These results support earlier surgical consideration and closer monitoring in women. Overall, the elastase+BAPN model effectively reproduces key clinical features of AAA, providing a useful platform for testing novel AAA therapies.

## Methods

### Animal Model

We induced AAAs in male (n = 27) and female (n = 7) C57Bl6/J mice aged an average of 13±1 weeks through a topical elastase surgical application and β-aminopropionitrile (BAPN) water-administration model. While image data from a subset of these were included in a previous publication (n = 15 males, n = 5 females; (18)), all data included here were analyzed independently and no analysis was repeated. Mice were anesthetized using 1-3% isoflurane in 0.6-1 L/minute of medical grade air. We opened the abdomen via a midline laparotomy and retracted the abdominal organs to achieve a clear view of the inferior vena cava and abdominal aorta. We cleared connective tissue from the ventral, left lateral, and dorsal sides of the vessel, leaving the aorta attached to the IVC. This created a small pocket in the connective tissue where we applied 5 µL of 5-10 mg/mL of porcine pancreatic elastase (Sigma Aldritch, Saint Louis, MO). The elastase was applied for 5 minutes and then the abdomen was flushed with saline three times. We closed the incision and administered 0.05 mL of long-acting buprenorphine as an analgesic. To counter ECM cross-linking repair, 0.2% BAPN (Sigma Aldritch), a lysyl oxidase inhibitor, was administered in the mouse drinking water starting two days prior to elastase application and continued throughout the study. Four male mice and three female mice recovered from surgery but died before the end of the study due to aortic rupture or surgical complications. These mice were excluded from further analysis, leaving the final study size n = 23 males and n = 4 females.

### Ultrasound Imaging

Two days prior to surgery, mice underwent baseline ultrasound imaging using a Vevo3100 high-frequency ultrasound system (FUJIFILM VisualSonics, transducer: MX550D (22-55 MHz)). Further imaging was collected at 7, 14, 21-, 28-, 42-, and 56-days post-surgery. At each imaging time point, axial brightness mode (B-mode) and E<u>C</u>G-gated kilohertz visualization (EKV), as well as sagittal B-mode, EKV, motion mode (M-mode), and color/pulsed wave Doppler images were collected. 3D and 4D ultrasound scans were also collected from the proximal aorta to the distal aorta towards the tail (step size=0.15 mm) at each timepoint. Maximum diameter measurements were collected from the 4D images and average peak systolic and end diastolic diameter at the largest portion of the vessel were collected from M-mode images using VevoLab analysis software (FUJIFILM VisualSonics). Green-Lagrange circumferential strain was calculated using systolic and diastolic diameters in the following formula (58): 3D segmentations of a subset of aortas were made using SimVascular (59). Initial estimates of the presence of thrombus were made from ultrasound images prior to confirming the presence of ILT using histology.

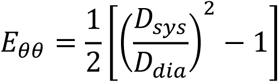

### Tissue Collection and Histology

After euthanasia, the aorta was extracted *in toto* starting at the left ventricular outflow tract and ending at the iliac arteries. Tissues were fixed in 4% paraformaldehyde for 48 hours and then stored in 70% ethanol prior to histopathologic processing. To prepare for histology, 1-mm thick sections of aorta were trimmed beginning at the level of the kidneys (n=10 sections/animal). The specimens were processed, embedded in paraffin, and stained according to standard histologic procedures. Each 1-mm section of aorta was stained with hematoxylin and eosin (H&E) and Movat’s Pentachrome (MPC). Microscopic slides were evaluated by a board-certified veterinary pathologist to characterize ILT, elastin disorganization, and the type of inflammation. A semi-quantitative histomorphological scale, based on standard histopathological morphology, was used that includes the amount of inflammatory infiltrate from H&E stained slides, and disorganization of elastin from MPC-stained slides. The histomorphologic scale was based on Table 1.

**Table 1:**
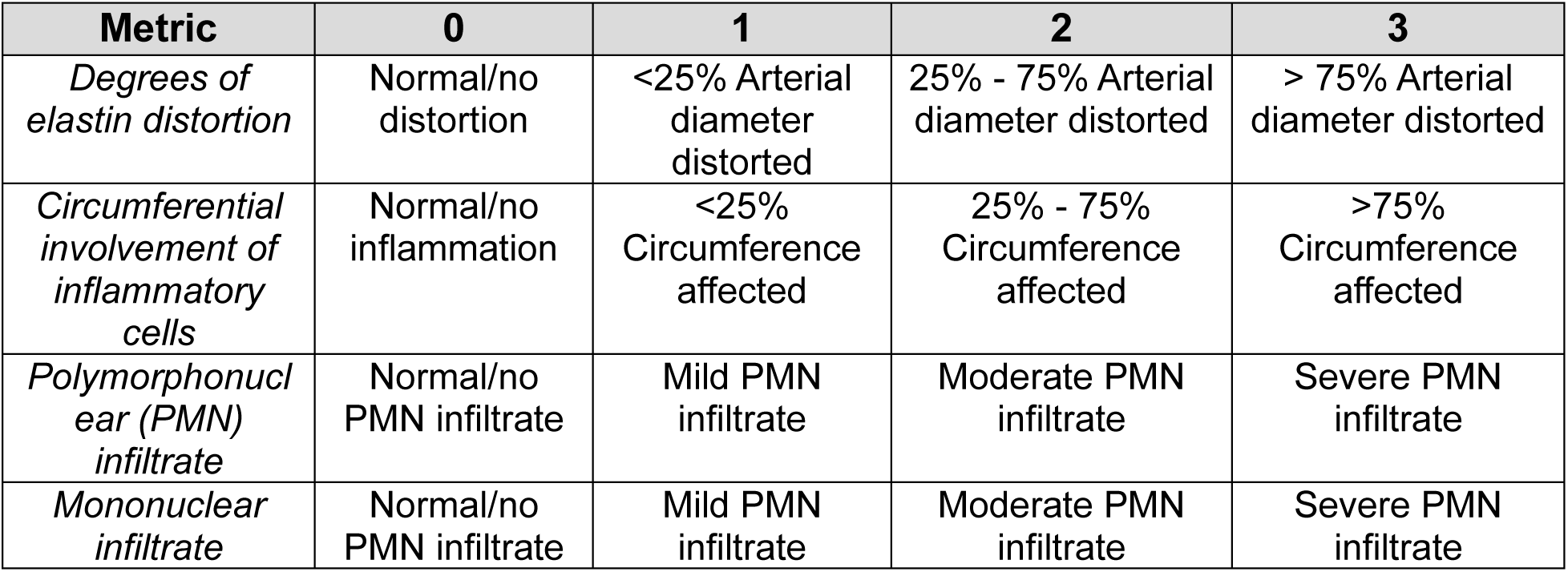
Semiquantitative Histological Analysis.

**Table 2:** Statistical Tests.

| Parameter | Normal Distribution | Statistical Test |
| --- | --- | --- |
| Combined Male Diameter | No | Kruskal-Wallis test with Dunn's multiple comparison test |
| Separated Male Diameter | Yes | Mixed-effects model with the Geisser-Greenhouse correction and Tukey's multiple comparison test |
| Combined Male Strain | Yes - log | Lognormal ordinary one-way ANOVA with Tukey's multiple comparison test |
| Separated Male Strain | Yes - log | Lognormal mixed-effects model with the Geisser-Greenhouse correction and Tukey's multiple comparison test |
| Tunica Intima Composition Males | Yes | Ordinary two-way ANOVA with Tukey's multiple comparison test |
| Tunica Media Composition Males | Yes | Ordinary two-way ANOVA with Tukey's multiple comparison test |
| Tunica Adventitia Composition Males | Yes - sqrt | Ordinary two-way ANOVA with a square root transformation and Tukey's multiple comparison test |
| Number of Intact Elastin Sheets Males | No | Kruskal-Wallis test with Dunn's multiple comparison test |
| Thickness of Elastin Sheets Males | Yes | Ordinary one-way ANOVA with Tukey's multiple comparison test |
| Diameter MvF | Yes | Mixed-effects model with the Geisser-Greenhouse correction and Tukey's multiple comparison test |
| Diameter MF ILT | Yes | Mixed-effects model with the Geisser-Greenhouse correction and Tukey's multiple comparison test |
| Number of Intact Elastin Sheets Females and Males | No | Mann-Whitney test |
| Number of Intact Elastin Sheets FM ILT | No | Kruskal-Wallis test with Dunn's multiple comparison test |
| Elastin Thickness MF | Yes | Welch's t-test |
| Elastin Thickness MF ILT | Yes | Ordinary one-way ANOVA with Tukey's multiple comparison test |
| Tunica Intima MF | Yes | Ordinary two-way ANOVA with Tukey's multiple comparison test |
| Tunica Intima MF ILT | Yes | Ordinary two-way ANOVA with Tukey's multiple comparison test |
| Tunica Media MF | Yes | Ordinary two-way ANOVA with Tukey's multiple comparison test |
| Tunica Media MF ILT | Yes | Ordinary two-way ANOVA with Tukey's multiple comparison test |
| Tunica Adventitia MF | Yes | Ordinary two-way ANOVA with Tukey's multiple comparison test |
| Tunica Adventitia MF ILT | Yes - sqrt | Ordinary two-way ANOVA with a square root transformation and Tukey's multiple comparison test |

Image Scope software was used to determine the aortic expansion from the aortic slice with the largest diameter (x64, Aperio). Images were collected at 20x zoom and separated into four randomly selected quadrants around the lumen. The thickest medial elastin sheet in each quadrant was quantified (PowerPoint), and manual counting of medial lamellar units was used to determine the number of distinct intact elastin sheets in the wall of the tunica media. Intimal hyperplasia, with dense and disordered elastin sheets, were not included in this analysis. A color segmentation tool (version 1.54p, ImageJ, NIH) was run to identify the extracellular matrix component composition of each layer of the aorta, finding the composition percentage of elastin, erythrocytes, collagen, fibrin, proteoglycans, and smooth muscle. One male mouse without detected ILT was excluded from histology analysis due to tissue processing issues. Color segmentation and elastin sheet number and thickness were also evaluated for n=3 aorta samples from a location higher than the aortic expansion to serve as controls.

### Scanning Electron Microscopy

A subset of fixed aorta slices on 1-mm were dehydrated using progressively increasing concentrations of ethanol and hexamethyldisilazane. After dehydration, samples were sputter coated with gold palladium. Low- and high-magnification images were collected in at least three locations for each sample.

### Statistics

All data was checked for normality using the Shapiro-Wilk’s test. Statistical tests for each parameter are listed in the table below. A *p* < 0.05 was considered significant.

## Supporting information

Combined Supplemental Files

