## Supplementary material for "Murine Abdominal Aortic Aneurysm Intraluminal Thrombus Composition and Structure": Combined Supplemental Files

Supplemental Table 1: *p*-values for Combined Male Diameter

| Dunn's multiple comparisons test | Summary | Adjusted P Value |
|---|---|---|
| Baseline vs. Day 7 | ns | >0.99 |
| Baseline vs. Day 14 | *** | <0.001 |
| Baseline vs. Day 21 | *** | <0.001 |
| Baseline vs. Day 28 | *** | <0.001 |
| Baseline vs. Day 42 | *** | <0.001 |
| Baseline vs. Day 56 | *** | <0.001 |
| Day 7 vs. Day 14 | ns | 0.49 |
| Day 7 vs. Day 21 | ** | 0.008 |
| Day 7 vs. Day 28 | *** | <0.001 |
| Day 7 vs. Day 42 | *** | <0.001 |
| Day 7 vs. Day 56 | *** | <0.001 |
| Day 14 vs. Day 21 | ns | >0.99 |
| Day 14 vs. Day 28 | ns | 0.51 |
| Day 14 vs. Day 42 | * | 0.03 |
| Day 14 vs. Day 56 | ** | 0.002 |
| Day 21 vs. Day 28 | ns | >0.99 |
| Day 21 vs. Day 42 | ns | >0.99 |
| Day 21 vs. Day 56 | ns | 0.19 |
| Day 28 vs. Day 42 | ns | >0.99 |
| Day 28 vs. Day 56 | ns | >0.99 |
| Day 42 vs. Day 56 | ns | >0.99 |

Supplemental Table 2: *p*-values for Separated Male Diameter

| Tukey's multiple comparisons test | | Summary | Adjusted P Value |
|---|---|---|---|
| Baseline | ILT vs. No ILT | ns | 0.83 |
| Day 7 | ILT vs. No ILT | ns | 0.05 |
| Day 14 | ILT vs. No ILT | * | 0.01 |
| Day 21 | ILT vs. No ILT | *** | <0.001 |
| Day 28 | ILT vs. No ILT | *** | <0.001 |
| Day 42 | ILT vs. No ILT | *** | <0.001 |
| Day 56 | ILT vs. No ILT | *** | <0.001 |
| ILT | Baseline vs. Day 7 | *** | <0.001 |
| | Baseline vs. Day 14 | *** | <0.001 |
| | Baseline vs. Day 21 | *** | <0.001 |
| | Baseline vs. Day 28 | *** | <0.001 |
| | Baseline vs. Day 42 | *** | <0.001 |
| | Baseline vs. Day 56 | *** | <0.001 |
| | Day 7 vs. Day 14 | ** | 0.008 |
| | Day 7 vs. Day 21 | *** | <0.001 |
| | Day 7 vs. Day 28 | *** | <0.001 |
| | Day 7 vs. Day 42 | *** | <0.001 |
| | Day 7 vs. Day 56 | *** | <0.001 |
| | Day 14 vs. Day 21 | ** | 0.002 |
| | Day 14 vs. Day 28 | *** | <0.001 |
| | Day 14 vs. Day 42 | *** | <0.001 |
| | Day 14 vs. Day 56 | *** | <0.001 |
| | Day 21 vs. Day 28 | ** | 0.001 |
| | Day 21 vs. Day 42 | *** | <0.001 |
| | Day 21 vs. Day 56 | *** | <0.001 |
| | Day 28 vs. Day 42 | ** | 0.003 |
| | Day 28 vs. Day 56 | *** | <0.001 |
| | Day 42 vs. Day 56 | * | 0.02 |
| No ILT | Baseline vs. Day 7 | * | 0.02 |
| | Baseline vs. Day 14 | *** | <0.001 |
| | Baseline vs. Day 21 | ** | 0.005 |
| | Baseline vs. Day 28 | ** | 0.005 |
| | Baseline vs. Day 42 | * | 0.01 |
| | Baseline vs. Day 56 | * | 0.01 |
| | Day 7 vs. Day 14 | * | 0.05 |
| | Day 7 vs. Day 21 | ns | 0.12 |
| | Day 7 vs. Day 28 | * | 0.04 |
| | Day 7 vs. Day 42 | * | 0.03 |
| | Day 7 vs. Day 56 | * | 0.04 |
| | Day 14 vs. Day 21 | ns | 0.24 |

| | | | |
|---|---|---|---|
| | Day 14 vs. Day 28 | * | 0.04 |
| | Day 14 vs. Day 42 | * | 0.04 |
| | Day 14 vs. Day 56 | * | 0.04 |
| | Day 21 vs. Day 28 | * | 0.03 |
| | Day 21 vs. Day 42 | * | 0.04 |
| | Day 21 vs. Day 56 | * | 0.04 |
| | Day 28 vs. Day 42 | ns | 0.09 |
| | Day 28 vs. Day 56 | ns | 0.05 |
| | Day 42 vs. Day 56 | ns | 0.06 |

Supplemental Table 3: *p*-values for Combined Male Strain

| Tukey's multiple comparisons test | Summary | Adjusted P Value |
|---|---|---|
| Baseline vs. Day 7 | ** | 0.007 |
| Baseline vs. Day 14 | *** | <0.001 |
| Baseline vs. Day 21 | *** | <0.001 |
| Baseline vs. Day 28 | *** | <0.001 |
| Baseline vs. Day 42 | *** | <0.001 |
| Baseline vs. Day 56 | *** | <0.001 |
| Day 7 vs. Day 14 | * | 0.04 |
| Day 7 vs. Day 21 | *** | <0.001 |
| Day 7 vs. Day 28 | *** | <0.001 |
| Day 7 vs. Day 42 | *** | <0.001 |
| Day 7 vs. Day 56 | *** | <0.001 |
| Day 14 vs. Day 21 | ns | 0.7 |
| Day 14 vs. Day 28 | ns | 0.69 |
| Day 14 vs. Day 42 | ns | 0.14 |
| Day 14 vs. Day 56 | ns | 0.71 |
| Day 21 vs. Day 28 | ns | >0.99 |
| Day 21 vs. Day 42 | ns | 0.96 |
| Day 21 vs. Day 56 | ns | >0.99 |
| Day 28 vs. Day 42 | ns | 0.95 |
| Day 28 vs. Day 56 | ns | >0.99 |
| Day 42 vs. Day 56 | ns | 0.95 |

Supplemental Table 4: *p*-values for Separated Male Strain

| Tukey's multiple comparisons test | | Summary | Adjusted P Value |
|---|---|---|---|
| Baseline | ILT vs. No ILT | ns | 0.4385 |
| Day 7 | ILT vs. No ILT | ns | 0.0506 |
| Day 14 | ILT vs. No ILT | * | 0.0342 |
| Day 21 | ILT vs. No ILT | ns | 0.0557 |
| Day 28 | ILT vs. No ILT | * | 0.0349 |
| Day 42 | ILT vs. No ILT | ns | 0.0556 |
| Day 56 | ILT vs. No ILT | ns | 0.3301 |
| ILT | Baseline vs. Day 7 | *** | 0.0005 |
| | Baseline vs. Day 14 | *** | 0.0003 |
| | Baseline vs. Day 21 | **** | <0.0001 |
| | Baseline vs. Day 28 | *** | 0.0007 |
| | Baseline vs. Day 42 | **** | <0.0001 |
| | Baseline vs. Day 56 | **** | <0.0001 |
| | Day 7 vs. Day 14 | ns | 0.2619 |
| | Day 7 vs. Day 21 | * | 0.0161 |
| | Day 7 vs. Day 28 | ns | 0.088 |
| | Day 7 vs. Day 42 | ** | 0.007 |
| | Day 7 vs. Day 56 | ns | 0.0926 |
| | Day 14 vs. Day 21 | ns | 0.9313 |
| | Day 14 vs. Day 28 | ns | 0.7986 |
| | Day 14 vs. Day 42 | ns | 0.576 |
| | Day 14 vs. Day 56 | ns | 0.9986 |
| | Day 21 vs. Day 28 | ns | 0.9985 |
| | Day 21 vs. Day 42 | ns | 0.9869 |
| | Day 21 vs. Day 56 | ns | 0.9447 |
| | Day 28 vs. Day 42 | ns | >0.9999 |
| | Day 28 vs. Day 56 | ns | 0.8165 |
| | Day 42 vs. Day 56 | ns | 0.3473 |
| No ILT | Baseline vs. Day 7 | ** | 0.0042 |
| | Baseline vs. Day 14 | *** | 0.0009 |
| | Baseline vs. Day 21 | ** | 0.0014 |
| | Baseline vs. Day 28 | **** | <0.0001 |
| | Baseline vs. Day 42 | *** | 0.0003 |
| | Baseline vs. Day 56 | **** | <0.0001 |
| | Day 7 vs. Day 14 | ns | 0.3562 |
| | Day 7 vs. Day 21 | * | 0.0241 |
| | Day 7 vs. Day 28 | * | 0.015 |
| | Day 7 vs. Day 42 | ** | 0.0031 |
| | Day 7 vs. Day 56 | ** | 0.0036 |
| | Day 14 vs. Day 21 | ns | 0.9705 |

| | | | |
|---|---|---|---|
| | Day 14 vs. Day 28 | ns | 0.885 |
| | Day 14 vs. Day 42 | ns | 0.5631 |
| | Day 14 vs. Day 56 | ns | 0.3426 |
| | Day 21 vs. Day 28 | ns | >0.9999 |
| | Day 21 vs. Day 42 | ns | 0.9944 |
| | Day 21 vs. Day 56 | ns | 0.9955 |
| | Day 28 vs. Day 42 | ns | 0.9779 |
| | Day 28 vs. Day 56 | ns | 0.9746 |
| | Day 42 vs. Day 56 | ns | >0.9999 |

Supplemental Table 5: Tunica Intima Composition Males

| Tukey's multiple comparisons test | | Summary | Adjusted P Value |
|---|---|---|---|
| Elastin | ILT vs. No ILT | ns | 0.536 |
| Cells | ILT vs. No ILT | ns | 0.393 |
| Collagen | ILT vs. No ILT | ns | 0.074 |
| Fibrin | ILT vs. No ILT | ns | 0.364 |
| Proteoglycans | ILT vs. No ILT | ns | 0.513 |
| ILT | Elastin vs. Cells | ns | 0.649 |
| | Elastin vs. Collagen | *** | <0.001 |
| | Elastin vs. Fibrin | ns | 0.86 |
| | Elastin vs. Proteoglycans | ns | >0.999 |
| | Cells vs. Collagen | * | 0.041 |
| | Cells vs. Fibrin | ns | 0.137 |
| | Cells vs. Proteoglycans | ns | 0.602 |
| | Collagen vs. Fibrin | **** | <0.001 |
| | Collagen vs. Proteoglycans | *** | <0.001 |
| | Fibrin vs. Proteoglycans | ns | 0.89 |
| No ILT | Elastin vs. Cells | ns | 0.07 |
| | Elastin vs. Collagen | ns | 0.062 |
| | Elastin vs. Fibrin | ns | 0.979 |
| | Elastin vs. Proteoglycans | ns | 0.778 |
| | Cells vs. Collagen | ns | >0.999 |
| | Cells vs. Fibrin | ns | 0.237 |
| | Cells vs. Proteoglycans | ns | 0.568 |
| | Collagen vs. Fibrin | ns | 0.217 |
| | Collagen vs. Proteoglycans | ns | 0.537 |
| | Fibrin vs. Proteoglycans | ns | 0.977 |

Supplemental Table 6: Tunica Media Composition Males

| Tukey's multiple comparisons test | | Summary | Adjusted P Value |
|---|---|---|---|
| Elastin | Control vs. No ILT | ns | 0.05 |
| | Control vs. ILT | * | 0.02 |
| | No ILT vs. ILT | ns | 0.85 |
| Cells | Control vs. No ILT | ns | 0.37 |
| | Control vs. ILT | ns | 0.07 |
| | No ILT vs. ILT | ns | 0.38 |
| Collagen | Control vs. No ILT | ns | 0.63 |
| | Control vs. ILT | ns | 0.22 |
| | No ILT vs. ILT | ns | 0.49 |
| Fibrin | Control vs. No ILT | ns | 0.95 |
| | Control vs. ILT | ns | 0.44 |
| | No ILT vs. ILT | ns | 0.35 |
| Proteoglycans | Control vs. No ILT | ** | 0.006 |
| | Control vs. ILT | *** | <0.001 |
| | No ILT vs. ILT | ns | 0.1 |
| Control | Elastin vs. Cells | * | 0.03 |
| | Elastin vs. Collagen | * | 0.03 |
| | Elastin vs. Fibrin | ns | 0.94 |
| | Elastin vs. Proteoglycans | ns | 0.09 |
| | Cells vs. Collagen | ns | >0.99 |
| | Cells vs. Fibrin | ** | 0.003 |
| | Cells vs. Proteoglycans | *** | <0.001 |
| | Collagen vs. Fibrin | ** | 0.002 |
| | Collagen vs. Proteoglycans | *** | <0.001 |
| | Fibrin vs. Proteoglycans | ns | 0.4 |
| No ILT | Elastin vs. Cells | *** | <0.001 |
| | Elastin vs. Collagen | *** | <0.001 |
| | Elastin vs. Fibrin | ns | 0.46 |
| | Elastin vs. Proteoglycans | ** | 0.005 |
| | Cells vs. Collagen | ** | 0.004 |
| | Cells vs. Fibrin | *** | <0.001 |
| | Cells vs. Proteoglycans | ** | 0.004 |
| | Collagen vs. Fibrin | *** | <0.001 |
| | Collagen vs. Proteoglycans | *** | <0.001 |
| | Fibrin vs. Proteoglycans | ns | 0.33 |
| ILT | Elastin vs. Cells | *** | <0.001 |
| | Elastin vs. Collagen | *** | <0.001 |
| | Elastin vs. Fibrin | ns | 0.93 |
| | Elastin vs. Proteoglycans | *** | <0.001 |
| | Cells vs. Collagen | *** | <0.001 |

| | | | |
|---|---|---|---|
| | Cells vs. Fibrin | *** | <0.001 |
| | Cells vs. Proteoglycans | ns | >0.99 |
| | Collagen vs. Fibrin | *** | <0.001 |
| | Collagen vs. Proteoglycans | *** | <0.001 |
| | Fibrin vs. Proteoglycans | *** | <0.001 |

Supplemental Table 7: Tunica Adventitia Composition Males

| Tukey's multiple comparisons test | | Summary | Adjusted P Value |
|---|---|---|---|
| Elastin | Control vs. No ILT | ns | 0.77 |
| | Control vs. ILT | ns | 0.41 |
| | No ILT vs. ILT | ns | 0.64 |
| Cells | Control vs. No ILT | ns | 0.65 |
| | Control vs. ILT | ns | 0.71 |
| | No ILT vs. ILT | ns | 0.99 |
| Collagen | Control vs. No ILT | ns | 0.95 |
| | Control vs. ILT | ns | >0.99 |
| | No ILT vs. ILT | ns | 0.8 |
| Fibrin | Control vs. No ILT | ns | 0.46 |
| | Control vs. ILT | ns | 0.86 |
| | No ILT vs. ILT | ns | 0.55 |
| Proteoglycans | Control vs. No ILT | ns | 0.34 |
| | Control vs. ILT | ns | 0.87 |
| | No ILT vs. ILT | ns | 0.29 |
| Control | Elastin vs. Cells | ** | 0.009 |
| | Elastin vs. Collagen | *** | <0.001 |
| | Elastin vs. Fibrin | ns | 0.75 |
| | Elastin vs. Proteoglycans | * | 0.05 |
| | Cells vs. Collagen | ** | 0.002 |
| | Cells vs. Fibrin | *** | <0.001 |
| | Cells vs. Proteoglycans | ns | >0.99 |
| | Collagen vs. Fibrin | *** | <0.001 |
| | Collagen vs. Proteoglycans | ** | 0.003 |
| | Fibrin vs. Proteoglycans | ** | 0.002 |
| No ILT | Elastin vs. Cells | *** | <0.001 |
| | Elastin vs. Collagen | *** | <0.001 |
| | Elastin vs. Fibrin | ns | 0.55 |
| | Elastin vs. Proteoglycans | ns | 0.2 |
| | Cells vs. Collagen | *** | <0.001 |
| | Cells vs. Fibrin | *** | <0.001 |
| | Cells vs. Proteoglycans | ns | 0.51 |
| | Collagen vs. Fibrin | *** | <0.001 |
| | Collagen vs. Proteoglycans | *** | <0.001 |
| | Fibrin vs. Proteoglycans | ** | 0.005 |
| ILT | Elastin vs. Cells | * | 0.01 |
| | Elastin vs. Collagen | *** | <0.001 |
| | Elastin vs. Fibrin | ** | 0.007 |
| | Elastin vs. Proteoglycans | * | 0.03 |
| | Cells vs. Collagen | *** | <0.001 |

| | | | |
|---|---|---|---|
| | Cells vs. Fibrin | *** | <0.001 |
| | Cells vs. Proteoglycans | ns | >0.99 |
| | Collagen vs. Fibrin | *** | <0.001 |
| | Collagen vs. Proteoglycans | *** | <0.001 |
| | Fibrin vs. Proteoglycans | *** | <0.001 |

Supplemental Table 8: Number of Intact Elastin Sheets Males

| Dunn's multiple comparisons test | Summary | Adjusted P Value |
|---|---|---|
| Control vs. No ILT | ns | 0.21 |
| Control vs. ILT | ** | 0.004 |
| No ILT vs. ILT | ns | 0.11 |

Supplemental Table 9: Thickness of Elastin Sheets Males

| Tukey's multiple comparisons test | Summary | Adjusted P Value |
|---|---|---|
| Control vs. No ILT | ns | 0.624 |
| Control vs. ILT | * | 0.047 |
| No ILT vs. ILT | ns | 0.057 |

Supplemental Table 10: Diameter Female vs. Male

| Tukey's multiple comparisons test | | Summary | Adjusted P Value |
|---|---|---|---|
| Baseline | Female vs. Male | ns | 0.06 |
| Day 7 | Female vs. Male | ns | 0.45 |
| Day 14 | Female vs. Male | ns | 0.44 |
| Day 21 | Female vs. Male | * | 0.03 |
| Day 28 | Female vs. Male | * | 0.03 |
| Day 42 | Female vs. Male | * | 0.02 |
| Day 56 | Female vs. Male | ns | 0.05 |
| Female | Baseline vs. Day 7 | ns | 0.27 |
| | Baseline vs. Day 14 | ns | 0.46 |
| | Baseline vs. Day 21 | * | 0.04 |
| | Baseline vs. Day 28 | * | 0.03 |
| | Baseline vs. Day 42 | * | 0.02 |
| | Baseline vs. Day 56 | ns | 0.05 |
| | Day 7 vs. Day 14 | ns | 0.67 |
| | Day 7 vs. Day 21 | * | 0.05 |
| | Day 7 vs. Day 28 | * | 0.02 |
| | Day 7 vs. Day 42 | ** | 0.004 |
| | Day 7 vs. Day 56 | ns | 0.06 |
| | Day 14 vs. Day 21 | ns | 0.39 |
| | Day 14 vs. Day 28 | ns | 0.31 |
| | Day 14 vs. Day 42 | ns | 0.11 |
| | Day 14 vs. Day 56 | * | 0.02 |
| | Day 21 vs. Day 28 | ns | 0.44 |
| | Day 21 vs. Day 42 | * | 0.04 |
| | Day 21 vs. Day 56 | ns | 0.1 |
| | Day 28 vs. Day 42 | * | 0.02 |
| | Day 28 vs. Day 56 | ns | 0.22 |
| | Day 42 vs. Day 56 | ns | 0.37 |
| Male | Baseline vs. Day 7 | *** | <0.001 |
| | Baseline vs. Day 14 | *** | <0.001 |
| | Baseline vs. Day 21 | *** | <0.001 |
| | Baseline vs. Day 28 | *** | <0.001 |
| | Baseline vs. Day 42 | *** | <0.001 |
| | Baseline vs. Day 56 | *** | <0.001 |
| | Day 7 vs. Day 14 | *** | <0.001 |
| | Day 7 vs. Day 21 | *** | <0.001 |
| | Day 7 vs. Day 28 | *** | <0.001 |
| | Day 7 vs. Day 42 | *** | <0.001 |
| | Day 7 vs. Day 56 | *** | <0.001 |
| | Day 14 vs. Day 21 | ** | 0.001 |

| | | | |
|---|---|---|---|
| | Day 14 vs. Day 28 | *** | <0.001 |
| | Day 14 vs. Day 42 | *** | <0.001 |
| | Day 14 vs. Day 56 | *** | <0.001 |
| | Day 21 vs. Day 28 | *** | <0.001 |
| | Day 21 vs. Day 42 | *** | <0.001 |
| | Day 21 vs. Day 56 | *** | <0.001 |
| | Day 28 vs. Day 42 | *** | <0.001 |
| | Day 28 vs. Day 56 | *** | <0.001 |
| | Day 42 vs. Day 56 | *** | <0.001 |

Supplemental Table 11: Diameter Female vs. Male ILT

| Tukey's multiple comparisons test | | Summary | Adjusted P Value |
|---|---|---|---|
| Baseline | Female ILT vs. Male No ILT | ns | 0.16 |
| Baseline | Female ILT vs. Male ILT | ns | 0.12 |
| Baseline | Male No ILT vs. Male ILT | ns | 0.97 |
| Day 7 | Female ILT vs. Male No ILT | ns | 0.47 |
| Day 7 | Female ILT vs. Male ILT | ns | 0.92 |
| Day 7 | Male No ILT vs. Male ILT | ns | 0.12 |
| Day 14 | Female ILT vs. Male No ILT | ns | 0.51 |
| Day 14 | Female ILT vs. Male ILT | ns | 0.88 |
| Day 14 | Male No ILT vs. Male ILT | * | 0.04 |
| Day 21 | Female ILT vs. Male No ILT | * | 0.02 |
| Day 21 | Female ILT vs. Male ILT | ns | 0.3 |
| Day 21 | Male No ILT vs. Male ILT | ** | 0.001 |
| Day 28 | Female ILT vs. Male No ILT | * | 0.01 |
| Day 28 | Female ILT vs. Male ILT | ns | 0.32 |
| Day 28 | Male No ILT vs. Male ILT | *** | <0.001 |
| Day 42 | Female ILT vs. Male No ILT | ** | 0.01 |
| Day 42 | Female ILT vs. Male ILT | ns | 0.35 |
| Day 42 | Male No ILT vs. Male ILT | ** | 0.001 |
| Day 56 | Female ILT vs. Male No ILT | * | 0.04 |
| Day 56 | Female ILT vs. Male ILT | ns | 0.28 |
| Day 56 | Male No ILT vs. Male ILT | ** | 0.003 |
| Female ILT | Baseline vs. Day 7 | ns | 0.27 |
| Female ILT | Baseline vs. Day 14 | ns | 0.46 |
| Female ILT | Baseline vs. Day 21 | * | 0.04 |
| Female ILT | Baseline vs. Day 28 | * | 0.03 |
| Female ILT | Baseline vs. Day 42 | * | 0.02 |
| Female ILT | Baseline vs. Day 56 | ns | 0.05 |
| Female ILT | Day 7 vs. Day 14 | ns | 0.67 |
| Female ILT | Day 7 vs. Day 21 | * | 0.05 |
| Female ILT | Day 7 vs. Day 28 | * | 0.02 |
| Female ILT | Day 7 vs. Day 42 | ** | 0.004 |
| Female ILT | Day 7 vs. Day 56 | ns | 0.06 |
| Female ILT | Day 14 vs. Day 21 | ns | 0.39 |
| Female ILT | Day 14 vs. Day 28 | ns | 0.31 |
| Female ILT | Day 14 vs. Day 42 | ns | 0.11 |
| Female ILT | Day 14 vs. Day 56 | * | 0.02 |
| Female ILT | Day 21 vs. Day 28 | ns | 0.44 |
| Female ILT | Day 21 vs. Day 42 | * | 0.04 |
| Female ILT | Day 21 vs. Day 56 | ns | 0.1 |
| Female ILT | Day 28 vs. Day 42 | * | 0.02 |

| | | | |
|---|---|---|---|
| | Day 28 vs. Day 56 | ns | 0.22 |
| | Day 42 vs. Day 56 | ns | 0.37 |
| Male No ILT | Baseline vs. Day 7 | * | 0.02 |
| | Baseline vs. Day 14 | *** | <0.001 |
| | Baseline vs. Day 21 | ** | 0.005 |
| | Baseline vs. Day 28 | ** | 0.005 |
| | Baseline vs. Day 42 | * | 0.01 |
| | Baseline vs. Day 56 | * | 0.01 |
| | Day 7 vs. Day 14 | * | 0.05 |
| | Day 7 vs. Day 21 | ns | 0.12 |
| | Day 7 vs. Day 28 | * | 0.04 |
| | Day 7 vs. Day 42 | * | 0.03 |
| | Day 7 vs. Day 56 | * | 0.04 |
| | Day 14 vs. Day 21 | ns | 0.24 |
| | Day 14 vs. Day 28 | * | 0.04 |
| | Day 14 vs. Day 42 | * | 0.04 |
| | Day 14 vs. Day 56 | * | 0.04 |
| | Day 21 vs. Day 28 | * | 0.03 |
| | Day 21 vs. Day 42 | * | 0.04 |
| | Day 21 vs. Day 56 | * | 0.04 |
| | Day 28 vs. Day 42 | ns | 0.09 |
| | Day 28 vs. Day 56 | ns | 0.05 |
| | Day 42 vs. Day 56 | ns | 0.06 |
| Male ILT | Baseline vs. Day 7 | *** | <0.001 |
| | Baseline vs. Day 14 | *** | <0.001 |
| | Baseline vs. Day 21 | *** | <0.001 |
| | Baseline vs. Day 28 | *** | <0.001 |
| | Baseline vs. Day 42 | *** | <0.001 |
| | Baseline vs. Day 56 | *** | <0.001 |
| | Day 7 vs. Day 14 | ** | 0.008 |
| | Day 7 vs. Day 21 | *** | <0.001 |
| | Day 7 vs. Day 28 | *** | <0.001 |
| | Day 7 vs. Day 42 | *** | <0.001 |
| | Day 7 vs. Day 56 | *** | <0.001 |
| | Day 14 vs. Day 21 | ** | 0.002 |
| | Day 14 vs. Day 28 | *** | <0.001 |
| | Day 14 vs. Day 42 | *** | <0.001 |
| | Day 14 vs. Day 56 | *** | <0.001 |
| | Day 21 vs. Day 28 | ** | 0.001 |
| | Day 21 vs. Day 42 | *** | <0.001 |
| | Day 21 vs. Day 56 | *** | <0.001 |
| | Day 28 vs. Day 42 | ** | 0.003 |

| | | | |
|---|---|---|---|
| | Day 28 vs. Day 56 | *** | <0.001 |
| | Day 42 vs. Day 56 | * | 0.02 |

Supplemental Table 12: Number of Intact Elastin Sheets Male and Female

| Mann Whitney test | |
|---|---|
| P value | 0.32 |
| Exact or approximate P value? | Exact |
| P value summary | ns |
| Significantly different (P < 0.05)? | No |
| One- or two-tailed P value? | Two-tailed |

Supplemental Table 13: Number of Intact Elastin Sheets Male and Female ILT

| Dunn's multiple comparisons test | Summary | Adjusted P Value |
|---|---|---|
| Male No ILT vs. Male ILT | * | 0.05 |
| Male No ILT vs. Female ILT | ns | 0.2 |
| Male ILT vs. Female ILT | ns | >0.99 |

Supplemental Table 14: Thickness of Elastin Sheets Male and Female

| Unpaired t test with Welch's correction | |
|---|---|
| P value | 0.4708 |
| P value summary | ns |
| Significantly different (P < 0.05)? | No |
| One- or two-tailed P value? | Two-tailed |
| Welch-corrected t, df | t=0.7857, df=4.571 |

Supplemental Table 15: Thickness of Elastin Sheets Male and Female ILT

| Tukey's multiple comparisons test | Summary | Adjusted P Value |
|---|---|---|
| Male No ILT vs. Male ILT | ns | 0.06 |
| Male No ILT vs. Female ILT | ns | 0.26 |
| Male ILT vs. Female ILT | ns | 0.99 |

Supplemental Table 16: Tunica Intima Composition Male and Female

| Tukey's multiple comparisons test | | Summary | Adjusted P Value |
|---|---|---|---|
| Elastin | Male vs. Female | ns | 0.063 |
| Cells | Male vs. Female | ns | 0.896 |
| Collagen | Male vs. Female | ns | 0.537 |
| Fibrin | Male vs. Female | ns | 0.564 |
| Proteoglycans | Male vs. Female | ns | 0.052 |
| Male | Elastin vs. Cells | * | 0.041 |
| | Elastin vs. Collagen | *** | <0.001 |
| | Elastin vs. Fibrin | ns | 0.996 |
| | Elastin vs. Proteoglycans | ns | 0.948 |
| | Cells vs. Collagen | ns | 0.187 |
| | Cells vs. Fibrin | * | 0.015 |
| | Cells vs. Proteoglycans | ns | 0.22 |
| | Collagen vs. Fibrin | *** | <0.001 |
| | Collagen vs. Proteoglycans | *** | <0.001 |
| | Fibrin vs. Proteoglycans | ns | 0.809 |
| Female | Elastin vs. Cells | ns | 0.072 |
| | Elastin vs. Collagen | *** | <0.001 |
| | Elastin vs. Fibrin | ns | 0.916 |
| | Elastin vs. Proteoglycans | * | 0.011 |
| | Cells vs. Collagen | ns | 0.523 |
| | Cells vs. Fibrin | ns | 0.389 |
| | Cells vs. Proteoglycans | ns | 0.96 |
| | Collagen vs. Fibrin | ** | 0.01 |
| | Collagen vs. Proteoglycans | ns | 0.902 |
| | Fibrin vs. Proteoglycans | ns | 0.107 |

Supplemental Table 17: Tunica Intima Composition Male and Female ILT

| Tukey's multiple comparisons test | | Summary | Adjusted P Value |
|---|---|---|---|
| Elastin | Male No ILT vs. Male ILT | ns | 0.8 |
| | Male No ILT vs. Female ILT | ns | 0.32 |
| | Male ILT vs. Female ILT | ns | 0.12 |
| Cells | Male No ILT vs. Male ILT | ns | 0.65 |
| | Male No ILT vs. Female ILT | ns | 0.88 |
| | Male ILT vs. Female ILT | ns | 0.98 |
| Collagen | Male No ILT vs. Male ILT | ns | 0.16 |
| | Male No ILT vs. Female ILT | ns | 0.38 |
| | Male ILT vs. Female ILT | ns | >0.99 |
| Fibrin | Male No ILT vs. Male ILT | ns | 0.62 |
| | Male No ILT vs. Female ILT | ns | 0.63 |
| | Male ILT vs. Female ILT | ns | 0.97 |
| Proteoglycans | Male No ILT vs. Male ILT | ns | 0.78 |
| | Male No ILT vs. Female ILT | ns | 0.28 |
| | Male ILT vs. Female ILT | ns | 0.1 |
| Male No ILT | Elastin vs. Cells | ns | 0.06 |
| | Elastin vs. Collagen | ns | 0.05 |
| | Elastin vs. Fibrin | ns | 0.98 |
| | Elastin vs. Proteoglycans | ns | 0.76 |
| | Cells vs. Collagen | ns | >0.99 |
| | Cells vs. Fibrin | ns | 0.21 |
| | Cells vs. Proteoglycans | ns | 0.54 |
| | Collagen vs. Fibrin | ns | 0.19 |
| | Collagen vs. Proteoglycans | ns | 0.51 |
| | Fibrin vs. Proteoglycans | ns | 0.97 |
| Male ILT | Elastin vs. Cells | ns | 0.63 |
| | Elastin vs. Collagen | *** | <0.001 |
| | Elastin vs. Fibrin | ns | 0.85 |
| | Elastin vs. Proteoglycans | ns | >0.99 |
| | Cells vs. Collagen | * | 0.03 |
| | Cells vs. Fibrin | ns | 0.12 |
| | Cells vs. Proteoglycans | ns | 0.58 |
| | Collagen vs. Fibrin | *** | <0.001 |
| | Collagen vs. Proteoglycans | *** | <0.001 |
| | Fibrin vs. Proteoglycans | ns | 0.88 |
| Female ILT | Elastin vs. Cells | ns | 0.07 |
| | Elastin vs. Collagen | *** | <0.001 |
| | Elastin vs. Fibrin | ns | 0.91 |
| | Elastin vs. Proteoglycans | * | 0.01 |
| | Cells vs. Collagen | ns | 0.52 |

| | | | |
|---|---|---|---|
| | Cells vs. Fibrin | ns | 0.38 |
| | Cells vs. Proteoglycans | ns | 0.96 |
| | Collagen vs. Fibrin | ** | 0.009 |
| | Collagen vs. Proteoglycans | ns | 0.9 |
| | Fibrin vs. Proteoglycans | ns | 0.1 |

Supplemental Table 18: Tunica Media Composition Male and Female

| Tukey's multiple comparisons test | | Summary | Adjusted P Value |
|---|---|---|---|
| Elastin | Male vs. Female | ns | 0.65 |
| Cells | Male vs. Female | ns | 0.8 |
| Collagen | Male vs. Female | * | 0.02 |
| Fibrin | Male vs. Female | ns | 0.81 |
| Proteoglycans | Male vs. Female | ** | 0.001 |
| Male | Elastin vs. Cells | *** | <0.001 |
| | Elastin vs. Collagen | *** | <0.001 |
| | Elastin vs. Fibrin | ns | 0.37 |
| | Elastin vs. Proteoglycans | *** | <0.001 |
| | Cells vs. Collagen | *** | <0.001 |
| | Cells vs. Fibrin | *** | <0.001 |
| | Cells vs. Proteoglycans | * | 0.04 |
| | Collagen vs. Fibrin | *** | <0.001 |
| | Collagen vs. Proteoglycans | *** | <0.001 |
| | Fibrin vs. Proteoglycans | *** | <0.001 |
| Female | Elastin vs. Cells | *** | <0.001 |
| | Elastin vs. Collagen | *** | <0.001 |
| | Elastin vs. Fibrin | ns | 0.88 |
| | Elastin vs. Proteoglycans | *** | <0.001 |
| | Cells vs. Collagen | ns | 0.65 |
| | Cells vs. Fibrin | ** | 0.006 |
| | Cells vs. Proteoglycans | ns | 0.53 |
| | Collagen vs. Fibrin | *** | <0.001 |
| | Collagen vs. Proteoglycans | ns | >0.99 |
| | Fibrin vs. Proteoglycans | *** | <0.001 |

Supplemental Table 19: Tunica Media Composition Male and Female ILT

| Tukey's multiple comparisons test | | Summary | Adjusted P Value |
|---|---|---|---|
| Elastin | Male No ILT vs. Male ILT | ns | 0.84 |
| | Male No ILT vs. Female ILT | ns | 0.8 |
| | Male ILT vs. Female ILT | ns | 0.97 |
| Cells | Male No ILT vs. Male ILT | ns | 0.34 |
| | Male No ILT vs. Female ILT | ns | 0.73 |
| | Male ILT vs. Female ILT | ns | 0.96 |
| Collagen | Male No ILT vs. Male ILT | ns | 0.44 |
| | Male No ILT vs. Female ILT | ns | 0.16 |
| | Male ILT vs. Female ILT | * | 0.02 |
| Fibrin | Male No ILT vs. Male ILT | ns | 0.3 |
| | Male No ILT vs. Female ILT | ns | 0.72 |
| | Male ILT vs. Female ILT | ns | 0.95 |
| Proteoglycans | Male No ILT vs. Male ILT | ns | 0.08 |
| | Male No ILT vs. Female ILT | *** | <0.001 |
| | Male ILT vs. Female ILT | * | 0.05 |
| Male No ILT | Elastin vs. Cells | *** | <0.001 |
| | Elastin vs. Collagen | *** | <0.001 |
| | Elastin vs. Fibrin | ns | 0.4 |
| | Elastin vs. Proteoglycans | ** | 0.002 |
| | Cells vs. Collagen | ** | 0.002 |
| | Cells vs. Fibrin | *** | <0.001 |
| | Cells vs. Proteoglycans | ** | 0.002 |
| | Collagen vs. Fibrin | *** | <0.001 |
| | Collagen vs. Proteoglycans | *** | <0.001 |
| | Fibrin vs. Proteoglycans | ns | 0.27 |
| Male ILT | Elastin vs. Cells | *** | <0.001 |
| | Elastin vs. Collagen | *** | <0.001 |
| | Elastin vs. Fibrin | ns | 0.91 |
| | Elastin vs. Proteoglycans | *** | <0.001 |
| | Cells vs. Collagen | *** | <0.001 |
| | Cells vs. Fibrin | *** | <0.001 |
| | Cells vs. Proteoglycans | ns | >0.99 |
| | Collagen vs. Fibrin | *** | <0.001 |
| | Collagen vs. Proteoglycans | *** | <0.001 |
| | Fibrin vs. Proteoglycans | *** | <0.001 |
| Female ILT | Elastin vs. Cells | *** | <0.001 |
| | Elastin vs. Collagen | *** | <0.001 |
| | Elastin vs. Fibrin | ns | 0.87 |
| | Elastin vs. Proteoglycans | *** | <0.001 |
| | Cells vs. Collagen | ns | 0.63 |

| | | | |
|---|---|---|---|
| | Cells vs. Fibrin | ** | 0.005 |
| | Cells vs. Proteoglycans | ns | 0.51 |
| | Collagen vs. Fibrin | *** | <0.001 |
| | Collagen vs. Proteoglycans | ns | >0.99 |
| | Fibrin vs. Proteoglycans | *** | <0.001 |

Supplemental Table 20: Tunica Adventitia Composition Male and Female

| Tukey's multiple comparisons test | | Summary | Adjusted P Value |
|---|---|---|---|
| Elastin | Male vs. Female | ns | 0.5 |
| Cells | Male vs. Female | * | 0.04 |
| Collagen | Male vs. Female | ns | 0.62 |
| Fibrin | Male vs. Female | ns | 0.78 |
| Proteoglycans | Male vs. Female | *** | <0.001 |
| Male | Elastin vs. Cells | *** | <0.001 |
| | Elastin vs. Collagen | *** | <0.001 |
| | Elastin vs. Fibrin | ns | 0.27 |
| | Elastin vs. Proteoglycans | ns | 0.49 |
| | Cells vs. Collagen | *** | <0.001 |
| | Cells vs. Fibrin | *** | <0.001 |
| | Cells vs. Proteoglycans | * | 0.04 |
| | Collagen vs. Fibrin | *** | <0.001 |
| | Collagen vs. Proteoglycans | *** | <0.001 |
| | Fibrin vs. Proteoglycans | ** | 0.004 |
| Female | Elastin vs. Cells | ns | 0.91 |
| | Elastin vs. Collagen | *** | <0.001 |
| | Elastin vs. Fibrin | ns | 0.98 |
| | Elastin vs. Proteoglycans | ** | 0.002 |
| | Cells vs. Collagen | *** | <0.001 |
| | Cells vs. Fibrin | ns | 0.62 |
| | Cells vs. Proteoglycans | * | 0.02 |
| | Collagen vs. Fibrin | *** | <0.001 |
| | Collagen vs. Proteoglycans | *** | <0.001 |
| | Fibrin vs. Proteoglycans | *** | <0.001 |

Supplemental Table 21: Tunica Adventitia Composition Male and Female ILT

| Tukey's multiple comparisons test | | Summary | Adjusted P Value |
|---|---|---|---|
| Elastin | Male No ILT vs. Male ILT | ns | 0.63 |
| | Male No ILT vs. Female ILT | ns | 0.66 |
| | Male ILT vs. Female ILT | ns | 0.28 |
| Cells | Male No ILT vs. Male ILT | ns | 0.99 |
| | Male No ILT vs. Female ILT | ns | 0.09 |
| | Male ILT vs. Female ILT | ns | 0.07 |
| Collagen | Male No ILT vs. Male ILT | ns | 0.79 |
| | Male No ILT vs. Female ILT | ns | 0.89 |
| | Male ILT vs. Female ILT | ns | >0.99 |
| Fibrin | Male No ILT vs. Male ILT | ns | 0.54 |
| | Male No ILT vs. Female ILT | ns | 0.71 |
| | Male ILT vs. Female ILT | ns | >0.99 |
| Proteoglycans | Male No ILT vs. Male ILT | ns | 0.28 |
| | Male No ILT vs. Female ILT | ** | 0.008 |
| | Male ILT vs. Female ILT | ns | 0.15 |
| Male No ILT | Elastin vs. Cells | *** | <0.001 |
| | Elastin vs. Collagen | *** | <0.001 |
| | Elastin vs. Fibrin | ns | 0.53 |
| | Elastin vs. Proteoglycans | ns | 0.19 |
| | Cells vs. Collagen | *** | <0.001 |
| | Cells vs. Fibrin | *** | <0.001 |
| | Cells vs. Proteoglycans | ns | 0.49 |
| | Collagen vs. Fibrin | *** | <0.001 |
| | Collagen vs. Proteoglycans | *** | <0.001 |
| | Fibrin vs. Proteoglycans | ** | 0.004 |
| Male ILT | Elastin vs. Cells | ** | 0.008 |
| | Elastin vs. Collagen | *** | <0.001 |
| | Elastin vs. Fibrin | ** | 0.005 |
| | Elastin vs. Proteoglycans | * | 0.02 |
| | Cells vs. Collagen | *** | <0.001 |
| | Cells vs. Fibrin | *** | <0.001 |
| | Cells vs. Proteoglycans | ns | >0.99 |
| | Collagen vs. Fibrin | *** | <0.001 |
| | Collagen vs. Proteoglycans | *** | <0.001 |
| | Fibrin vs. Proteoglycans | *** | <0.001 |
| Female ILT | Elastin vs. Cells | ns | 0.57 |
| | Elastin vs. Collagen | *** | <0.001 |
| | Elastin vs. Fibrin | ns | 0.9 |
| | Elastin vs. Proteoglycans | *** | <0.001 |
| | Cells vs. Collagen | *** | <0.001 |

| | | | |
|---|---|---|---|
| | Cells vs. Fibrin | ns | 0.13 |
| | Cells vs. Proteoglycans | * | 0.01 |
| | Collagen vs. Fibrin | *** | <0.001 |
| | Collagen vs. Proteoglycans | ** | 0.003 |
| | Fibrin vs. Proteoglycans | *** | <0.001 |

Supplemental Figure 1: Female and male mice with and without ILT

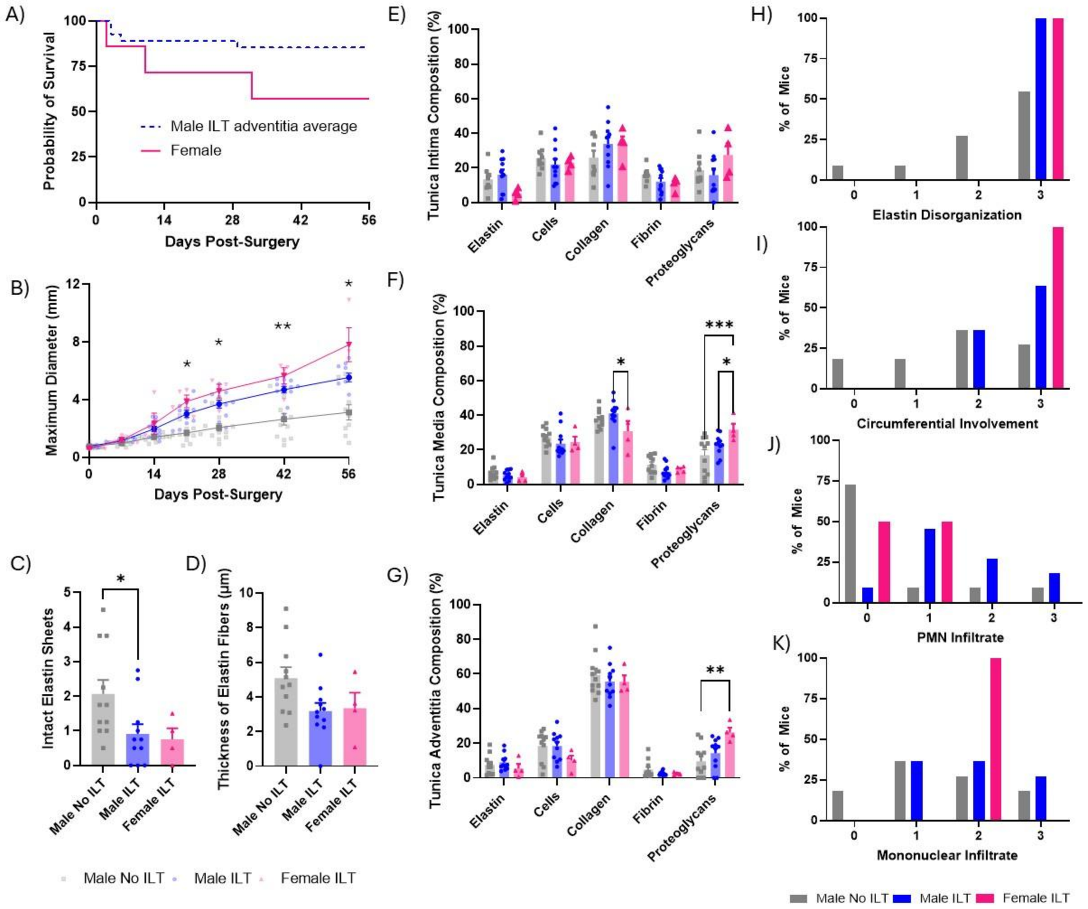


*S1: Female mice demonstrated greater response to elastase-BAPN procedure compared to males. A) Number of surviving male and female mice. Final numbers of males included n = 12 ILT, n = 11 no ILT, with a mortality of n = 4 after surgery. Final numbers of females included n = 4 ILT, no = 0 no ILT, with a mortality of n = 3 after surgery. B) Diameter of all samples split into female ILT (pink), male ILT (blue), and male no ILT (grey) groups. Bolded data points presented as averages±SE each timepoint. Statistical significance calculated via a mixed-effects model with the Geisser-Greenhouse correction and Tukey's multiple comparison test. *p<0.05;**p<0.01 - statistically significant difference between difference between female ILT and male no ILT groups. n = 12 male no ILT; n = 11 male ILT; n = 4 female ILT. C) Comparison of the number of intact elastin sheets between males with no ILT, males with ILT, and females with ILT, achieved via manual counting of the intact elastin sheets in the wall of the aorta. Statistical significance calculated via a Kruskal-Wallis test with Dunn's multiple comparison test. D) Comparison of the thickness of the elastin lamellar units between males with no ILT, males with ILT, and females with ILT. Statistical significance calculated via an ordinary one-way ANOVA with Tukey's multiple comparison test. E) tunica adventitia, F) tunica media, and G) tunica intima for each Movat's pentachrome stained tissue sample were quantified using ImageJ color segmentation. Statistical significance calculated via an ordinary two-way ANOVA with Tukey's multiple comparison test. Data from tunica adventitia composition underwent a square root transformation to address normality. *p<0.05;**p<0.01; ***p<0.001. Bars presented as averages±SE each timepoint. Semi-quantitative assessments of H) elastin disorganization where 0 is normal/no distortion; 1 is less than 25%; 2 is between 25% and 75%; and 3 is greater than 75% of the aortic circumference is affected; I) circumferential involvement of inflammatory cells (% area affected inflammation) where 0 is normal/no inflammation; 1 is less than 25%; 2 is between 25% and 75%; and 3 is greater than 75% of the aortic circumference affected; J) polymorphonuclear infiltrate (acute inflammation) where 0 is normal/no polymorphonuclear infiltrate and 3 is severe polymorphonuclear infiltrate; and K) mononuclear infiltrate (chronic inflammation) where 0 represents normal/no mononuclear infiltrate and 3 is severe mononuclear infiltrate. Semi-quantitative scoring determined by a board-certified veterinary histopathologist. Intima: n=9 male no ILT; n = 11 male ILT; n=4 female; fewer samples were evaluated of the tunica intima due to damage and obscuring of the intima seen in some of the non-thrombus forming samples. All other groups n=11 male no ILT; n = 11 male ILT; n=4 female ILT.*
